# Microhabitat acclimatization shifts physiological baselines and thermal tolerance of the symbiotic anemone, *Anthopleura elegantissima*

**DOI:** 10.1101/2023.05.15.540861

**Authors:** Maria Ruggeri, Wyatt C Million, Lindsey Hamilton, Carly D Kenkel

## Abstract

Contemporary organisms in extreme environments can give insight into how species will respond to environmental change. The intertidal forms an environmental gradient where stress increases with tidal height. Here, we explore the contribution of fixed genotypic and plastic environmental effects on thermal tolerance of the intertidal anemone *Anthopleura elegantissima* and its algal symbionts using a laboratory-based tank experiment. High intertidal anemones had lower baseline symbiont-to-host cell ratios under control conditions, but their symbionts had higher baseline maximum quantum yield compared to low intertidal anemone symbionts, despite identical symbiont communities. High intertidal anemones maintained greater maximum quantum yield and symbiont-to-host cell ratios under heat stress compared to low intertidal anemones, suggesting that high intertidal holobionts have greater thermal tolerance. However, thermal tolerance of clonal anemones acclimatized to different zones was not explained by tidal height alone, indicating emersion duration is not the sole environmental driver of physiological variation. Fixed genotypic effects also influenced physiological baselines, but did not modulate thermal tolerance, demonstrating thermal tolerance is largely driven by environmental history. These results indicate that this symbiosis is highly plastic and may be able to rapidly acclimatize to climate change, defying the convention that symbiotic organisms are more susceptible to environmental stress.

## Introduction

The ability to tolerate environmental extremes can be driven by fixed or plastic effects, but both play a role in species persistence over ecological and evolutionary timescales (Kumarathunge et al., 2019; Palumbi et al., 2014; Somero, 2010). Fixed genetic effects determine standing trait variation and tolerance limits between individuals, populations, and species, whereas plastic effects shift trait values within an individual. Fixed genetic effects can drive population-level adaptation over generations, whereas plastic effects can influence survival of an individual, potentially facilitating adaptation over longer time-scales (Kelly, 2019). At small spatial scales, phenotypic differentiation is likely due to physiological plasticity, or acclimatization, as high gene flow can limit adaptation (Lenormand, 2002; Richardson et al., 2014; Sultan & Spencer, 2002). However, steep environmental gradients can increase the strength of selection over high gene flow, which promotes adaptive divergence over microgeographic scales, within a population’s dispersal neighborhood (Felsenstein, 1976; Hendry et al., 2001; Richardson et al., 2014). It is therefore important to understand the patterns of and mechanisms facilitating stress tolerance in contemporary populations inhabiting extreme environments to inform predictions of how species will respond to climate extremes expected during global climate change.

Marine populations are generally well connected due to high dispersal, yet microgeographic trait divergence has been documented, indicating the existence of strong selection gradients over small spatial scales (Sanford & Kelly, 2011). The intertidal zone, in particular, forms a steep environmental gradient that has been shown to outweigh the homogenizing effect of gene flow (Sanford & Kelly, 2011; Somero, 2010). For example, congeneric species of crabs and snails exhibit divergence in thermal tolerance traits within meters of intertidal height, reflecting the vertical zonation of these species and implicating temperature as a potential driver of adaptation and speciation (Stillman & Somero, 2000; Tomanek & Somero, 1999). However, few studies have examined intraspecific variation in thermal tolerance traits across the intertidal. A common garden experiment using F1 progeny of *Crassostrea gigas* found higher survival, metabolic rates, and gene expression plasticity in intertidal-origin oysters compared to subtidal populations, indicating adaptive divergence across microgeographic scales (Li et al., 2018). However, a reciprocal transplant in *Mytilus californianus* found that juvenile mussels acclimatized to greater thermal variability increased their thermal tolerance regardless of tidal height origin, suggesting plasticity can also drive thermal tolerance divergence across the intertidal (Gleason et al., 2018). Therefore, the relative roles of adaptation versus plasticity in driving thermal tolerance variation across microgeographic scales remains unclear.

The importance of fixed and plastic effects at microgeographic scales can also be influenced by symbiotic associations, especially when partners differ in their capacity for gene flow (Thornhill et al., 2017) or when physiological thresholds are determined by holobiont rather than individual partner limits (Rosenberg et al., 2007; Zilber-Rosenberg & Rosenberg, 2008). Tropical corals are a particularly thermally sensitive symbiosis, where dysbiosis, or dissociation of dinoflagellate symbionts from the host cnidarian commonly known as thermal bleaching, occurs only 1-2°C above average maximum monthly temperatures (Hoegh-Guldberg, 1999). Yet thermal tolerance limits can differ over microgeographic scales. Previous research in acroporid corals from adjacent backreef pools 500 meters apart with contrasting daily thermal variability caused by tidal inundations has indicated a role for both fixed and plastic effects in driving divergence of thermal tolerance traits in the holobiont (reviewed in Thomas et al., 2018). However, another study in poritid corals from the same sites found that corals from the highly variable environment did not exhibit significant differences in thermal tolerance compared to those from less variable environments (Klepac & Barshis, 2020), complicating the idea that more thermally variable environments always increase thermal tolerance and highlighting the importance of understanding the mechanisms that drive intraspecific variation in thermal tolerance across small spatial scales.

Populations of the symbiotic anemone, *Anthopleura elegantissima*, are distributed throughout the intertidal zone along the Pacific coast of North America where it experiences extreme environmental fluctuations in temperature, light, food and water availability (Bingham et al., 2011; Dimond et al., 2011). Similar to reef-building corals, these anemones form a mutualistic symbiosis with the dinoflagellate, *Breviolum muscatinei* (formerly Symbiodinium, ITS2 type B4, Lajeunesse and Trench, 2000; LaJeunesse et al., 2018). Despite the general thermal sensitivity of cnidarian-dinoflagellate symbioses, these anemones can experience temperature fluctuations of up to 20°C during tidal exposure (Bingham et al., 2011). In addition to sexual reproduction, *A. elegantissima* also reproduce asexually to form large clonal aggregations, allowing the opportunity to leverage genetically identical individuals to tease apart plasticity from host genotypic effects. Anemones in the center of aggregations experience reduced thermal fluctuations compared to clonemates on the outer edges (Bingham et al., 2011), which could lead to differences in thermal tolerance within clonal anemones of a single genotype. This provides a unique system in which to explore the mechanisms leading to exceptional holobiont thermal tolerance across small spatial scales.

In this study, we explored the contribution of fixed and plastic effects on holobiont thermal tolerance of *A. elegantissima* acclimatized to different thermal environments. Replicate anemone aggregations were sampled from high and low intertidal environments, including clonal anemones taken from the center and edge of each aggregation, and exposed to control or elevated temperature (+10°C) for 10 days (Fig 1). Thermal tolerance was assessed by measuring maximum quantum yield of PSII, as an indicator of photodamage to the symbiont, and bleaching phenotypes, including symbiont to host cell ratios and chlorophyll concentration. 2bRAD genotyping was performed to verify host clonality within and among aggregations (Manzello et al., 2019; Wang et al., 2012) and symbiont community composition was assessed using amplicon sequencing of the ITS2 locus (Hume et al., 2019). Genotyping revealed that some independent aggregations spanning multiple tidal heights were found to be clonal, which provided an opportunity to explore thermal tolerance plasticity across the intertidal. We found that physiological baselines were defined by both microhabitat and host genotype, but microhabitat was the primary driver of thermal tolerance variation, suggesting acclimatization can greatly increase holobiont tolerance.

**Figure 1:**
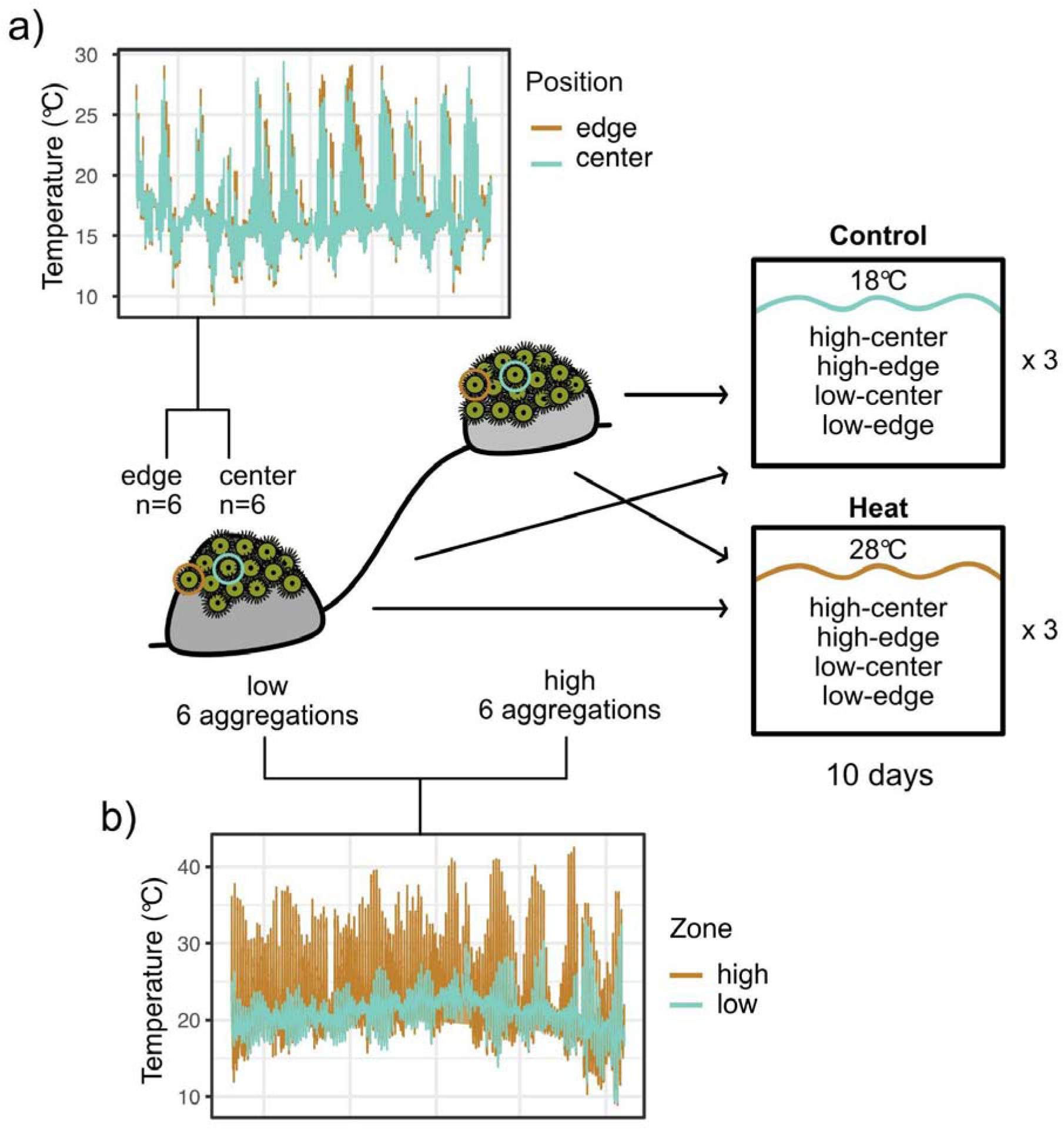
Diagram of sampling, experimental design, and thermal characteristics of sampling locations. Temperature profiles contrasting the center (blue) versus the edge (orange) of a low intertidal aggregation are shown in the top left (a) and temperature profiles contrasting the low (blue) versus high (orange) intertidal are shown in the bottom left (b).

## Methods

### Environmental characterization

Onset HOBO Tidbit MX400 temperature loggers were deployed *in-situ* in the high (+1.37m) and low (+0.47m) intertidal at Shark Harbor (latitude 33° 23’ 0.8226”, longitude −118° 28’ 24.0594”) from November 2019-November 2022 to characterize the range of environmental conditions experienced within each microhabitat (Fig. 1a). Temperature was recorded every 5 minutes and Bluetooth Off Water Detect was used to record when loggers were exposed to air. HOBO pendant loggers were also deployed in the center and edge of one low intertidal aggregation to characterize temperature variation within an aggregation (Fig. 1b). Due to difficulties in logger placement, no logger was able to be placed in the center of the high aggregation. Therefore temperature data from the center and edge of a high aggregation at a nearby site in Los Angeles County (Point Dume: latitude 34° 0’ 7.5486”, longitude −118° 48’ 18.1044”) were also collected.

### Heat stress experiment

Anemones were collected from Shark Harbor on Catalina Island, CA on November 13^th^, 2020 during low tide under a California Department of Fish and Wildlife specific-use scientific collecting permit (S-183050013-18305-001). To capture an environmental gradient, 6 distinct aggregations were sampled from the high intertidal and 6 distinct aggregations from the low intertidal using a micro-spatula (Fig. 1b). Aggregations were marked with epoxy and absolute tidal height was later measured with a DeWalt rotary laser level in November 2022 (Table S1). Low intertidal anemones were sampled from an average tide height of 0.363m (range 0.233m-0.783, Fig S1) and high intertidal anemones were sampled from an average tide height of 1.035m (range 0.834m-1.193m, Fig S1). Low aggregation 3 was partially buried during tidal height sampling and actual sampling height was likely lower than +0.783m. From each aggregation, 6 anemones were also collected from the inside (center) and 6 from the outer margins of the aggregation (edge), totaling 144 samples (Fig 1a). Anemones were placed in plastic bags full of seawater and transported to the Wrigley Marine Science Center, where they were cleaned of all debris, blotted dry, and weighed prior to being settled onto aragonite plugs set inside 29 mL plastic cups (Fig S2). Cups were covered with mesh for 24 hours to ensure attachment and distributed into six aquaria, equipped with Marine LED lights (150 umol/photons/m^2^) and submersible pumps, so that each tank contained one replicate of each aggregation-position combination (Fig 1). Aquaria were submerged in water baths in a temperature controlled room at 18°C and anemones were acclimated to these control conditions for 7 days prior to experimentation. Pulse-amplitude modulated fluorometry (PAM) was used daily to monitor photo-acclimation. Stablization of values, indicating photoacclimation, was observed on day 5 (Fig S3).

On day 8, temperatures in treatment tanks (n=3) were increased at 1°C per hour using 100-Watt heaters (Aquaneat) until they reached 28°C and were held at that temperature for 10 days (28.1± 0.239°C on average), while the control tanks (n=3) remained at 18°C (18.3 ± 0.223°C on average, Fig S4). Anemones were fed frozen brine shrimp twice weekly and 25% water changes were performed after feeding. Salinity was measured daily using a refractometer and averaged 35 ppt. At the end of the experimental period, anemones were relaxed in a 1:1 seawater: 0.3M MgCl_2_ solution and tentacle biopsies were taken from each individual and immediately frozen at −80°C.

### Thermal physiology

Maximum quantum yield (MQY) of photosystem II was measured daily during the experimental period following one hour of dark acclimation to assess photosynthetic performance using a Walz diving-PAM with the following settings: measuring intensity = 6, saturation intensity = 6, pulse width = 0.8, gain = 6, damp = 2. Contraction was induced by gently prodding anemones and MQY measurements were taken of the body column.

Frozen tentacle biopsies were thawed and homogenized in 1 mL extraction buffer (100 mM Tris HCl, 0.05 mM DTT, pH 7.8) using a rotor stator (VWR 200 homogenizer). Host and symbiont cells were then separated via centrifugation for 3 min at 1500 x G. Resulting host supernatant was removed and stored in 2 mL cell culture plates at −80°C and symbiont pellets were stored in 1.5 mL tubes at −80°C until protein and chlorophyll quantification, respectively. Protein concentration was quantified from the host supernatant using a BCA protein assay kit and normalized by the volume of elution buffer added during homogenization. To extract chlorophyll, symbiont pellets were resuspended in 90% acetone with 5-6 1mm stainless steel beads (McMaster-Carr) and homogenized in an Omni bead beater (30s at 4m/s for 3 cycles) to lyse cell walls. Chlorophyll samples were incubated at −20°C overnight before measuring the absorbance at 630, 647, and 664nm in triplicate using a Bio-tek, Synergy, H1M Microplate reader. Chlorophyll a was calculated from absorbance values using the equation from Ritchie (2008) and normalized by host soluble protein content.

### Symbiont to host cell ratios

DNA was extracted from frozen tentacles following Wayne’s method (Wilson et al., 2002), using an enzymatic lysis followed by alcohol precipitation. DNA was not obtained from eight samples due to failed extractions, resulting in a final sample size of 136. Symbiont to host cell ratios were quantified via qPCR by amplifying a host-specific ATPase gene and a symbiont-specific cp23S-rDNA gene (Dziedzic, 2019, Dziedzic et al., *in prep*). Primer-specific reactions were run in duplicate on an Agilent AriaMx with 2x Brilliant III Ultra-Fast SYBR MM, 1 mM reference dye, 400 nM forward and reverse primer, and 20 ng of template DNA in 20 ul reactions. Primer-specific Ct values of technical replicates were averaged after reference dye correction and symbiont to host ratios were calculated as the fold change of symbiont to host Ct multiplied by 2 to correct for copy number [(2^(Ct_host_ – Ct_symbiont_))*2] following (Cunning & Baker, 2012).

### Host genotyping

To verify clonality of the host, a subset of 48 DNA samples representing each aggregation-position-treatment combination were genotyped using 2bRAD (Wang et al., 2012) using a reduced representation approach targeting 1/16^th^ of BcgI sites. 2bRAD libraries were then sequenced on the NextSeq 500 by the USC Molecular Genomics core in June 2022. Two sequencing runs were performed yielding a total of 26.6 M raw reads.

Bioinformatic analysis was performed on the USC CARC HPC system following the pipeline described in https://github.com/z0on/2bRAD_denovo. First, reads were de-multiplexed based on internal ligation adaptor barcodes prior to adaptor trimming and removal of PCR duplicates using a custom perl script. Reads were then quality filtered to retain only 99% accurate base calls over 100% of the read using the fastx-toolkit and reads containing library adaptor sequences were removed. Following de-multiplexing, quality filtering and removal of PCR duplicates, 133,597 reads per sample remained from run 1 and 115,373 reads from run 2, totaling 248,170 reads per sample on average. High quality reads were concatenated across samples and competitively mapped to a combined *A. elegantissima* genome (Elder, 2020) and *Breviolum minutum* (formerly *Symbiodinium minutum*, ITS2 type B1) draft genome (Shoguchi et al., 2013) using Bowtie2 (Langmead & Salzberg 2012).

ANGSD 0.933 (Korneliussen et al., 2014) was used to call genotype likelihoods for reads exhibiting high quality matches to the anemone genome after filtering for high confidence SNPs present in at least 75% of samples. Hierarchical clustering of the resulting identity by state (IBS) matrix was used to identify clonal individuals based on known technical replicates as described in (Manzello et al., 2019). The genotype designation for one sample (11-O-1) was unclear, as this sample clustered with genet 13, while other replicates from this aggregation clustered with genet 10 (Fig S5). We therefore decided to re-prepare this sample along with the remaining replicates of aggregations 11 and 13, which were sequenced on the NextSeq 2000 by the USC Molecular Genomics core in November 2022 resulting in 17.4 M raw reads. Samples were processed and analyzed using the pipeline described above, which confirmed anemones from aggregation 11 were unique from aggregation 13 (Fig S6).

### Symbiont community profiling

Amplicon libraries of the ITS2 region were prepared using primers designed by Hume et al (2019). PCR reactions were prepared with 50 ng of input DNA and amplified using the following profile [98°C 0:30, (98°C 0:10 | 56°C 1:00 | 72°C 0:30) x 22, 72°C 5:00]. A second PCR was then run to incorporate sample-specific barcodes [98°C 0:30, (98°C 0:10 | 59°C 0:30 | 72°C 0:30) x 6, 72°C 2:00] and products were visualized by gel electrophoresis. 12-sample pools were combined based on band intensity, which were quantified spectrophotometrically using the Quant iT PicoGreen dsDNA assay kit. Equal amounts of each pool was added to the final library. Paired-end 250 bp reads were sequenced on the MiSeq v2 by the USC Molecular Genomics core with 30% PhiX spike-in yielding a read depth of ~29,000 reads per sample across two runs (nano and v2 chemistry) in June and July 2022. Forward and reverse reads were concatenated across runs and analyzed in SymPortal using default settings. ITS2 profiles were generated in SymPortal by collapsing co-occuring defining intragenomic variants (DIVs) within samples (Hume et al., 2019).

### Statistical Analyses

Statistical analyses were performed in R v4.2.1. Differences in environmental parameters (temperature and exposure) were evaluated across intertidal zones and positions within aggregations using Welch’s t-test assuming unequal variances across groups. All traits and their residuals were screened for outliers and normality and transformations were applied where appropriate. A log transformation was applied to both symbiont to host cell ratios and anemone weight. For the full dataset, linear mixed models were used to evaluate the effects of treatment, tidal height, position within aggregation, and their interaction on physiological traits (MQY, symbiont to host cell ratios, chlorophyll), with random effects of tank and host genotype using the lme4 package (Bates et al., 2015). Tidal height was evaluated both as a continuous metric as well as binned by intertidal zone (i.e. high vs low). Binning low aggregation 3 (+0.783m) with the high intertidal aggregations (Table S2) or using continuous tidal height measurements (Table S3) did not change the overall model results, therefore we have elected to present results binned by original tidal zone designations (Table S4). Where there was a significant random effect of host genotype, an additional model was run with host genotype as an independent fixed effect and as an interaction term with treatment to facilitate pairwise comparisons across genets using the ‘emmeans’ package in R (Lenth, 2022). Linear regressions between traits (MQY, symbiont to host cell ratios, and chlorophyll) were evaluated using linear models to explore the relationship between commonly measured bleaching phenotypes. As weight was only measured initially, treatment and tank were not included in models for anemone weight.

2bRAD genotyping revealed multiple aggregations across tidal heights belonging to a single genotype (Fig S5, Fig S1). These multi-aggregation genets, 1 and 10, were subset to explore plastic trait variation within genotypes. Within each genet, linear mixed models were run to evaluate the fixed effects of aggregation, treatment and their interaction on physiological traits, with tank as a random effect. Statistical scripts and input files can be found at https://github.com/mruggeri55/AeleHeatStress.

## Results

### Microhabitat gradients

The high intertidal (+1.37m) experienced warmer temperatures on average (+1.2°C) compared to the low intertidal (+0.47m) in the summer through fall of 2022 (high 22.1°C; low 20.9°C, p<0.001, Fig 1). The high intertidal also had 9.76°C greater daily thermal range on average compared to the low intertidal during this same period (high 16.1°C, low 6.34°C, p<0.001, Fig 1) and reached a maximum temperature of 42.6°C compared to 33°C in the low intertidal. Average temperature did not differ between center and edge anemones in the low intertidal at Shark Harbor (center 16.5°C, edge 16.5°C, p=0.64), but did significantly differ in the high intertidal at Point Dume, where edge anemones experienced about 0.1°C greater temperature on average (center 15.1°C, edge 15.2°C, p=0.002, Fig S7). Although not significantly different, daily thermal range was on average 0.46°C greater on the edge of aggregations versus the center in the low intertidal (center 5.76°C, edge 6.22°C, p=0.23, Fig 1). However, daily thermal range within aggregations did significantly differ at Point Dume, with 0.94°C greater daily thermal range on the edge versus center of a high intertidal aggregation (center 5.53°C, edge 6.47°C, p=0.001, Fig S7), and high intertidal edges reaching a maximum temperature 3°C above the center of the aggregation (center 26°C, edge 29°C, Fig S7).

### Shifting physiological baselines among intertidal zones and host genotypes

All genotyped samples hosted an identical symbiont profile (1379_B/1393_B-1607_B-B4-1608_B-1418_B-1612_B) corresponding to the species *Breviolum muscatinei* (LaJeunesse & Trench 2000; blastn 98.51% identity, E=5e^-129^, Fig S8), previously identified as the dominant associate of *A. elegantissima* in southern California (Sanders & Palumbi, 2011; Secord & Augustine, 2005).

Among aggregations, low intertidal anemones tended to have higher symbiont to host cell ratios, but their symbionts had reduced maximum quantum yield in comparison to their higher intertidal counterparts (Fig 2). This was supported by a significant fixed effect of intertidal zone on MQY (p=0.015, Table S4). Although there was no fixed effect of intertidal zone on symbiont to host cell ratios (p=0.348, Table S4), MQY (p<0.001) was a significant predictor of symbiont to host cell ratios and varied by tidal zone (p=0.011) (Fig 2, Table S5), indicating that symbiont performance differs among anemone aggregations inhabiting different intertidal zones and may influence symbiont density. Chlorophyll a was weakly correlated with both MQY (p<0.01, R^2^=0.06, Fig S9a) and symbiont to host cell ratios (p<0.001, R^2^=0.17, Fig S9b), but there was no effect of intertidal zone. Position within an aggregation did not affect MQY, symbiont to host cell ratios, or chlorophyll concentration (Fig S10, Table S4). However, position did have a significant effect on weight (p<0.001), where clonemates in the center of aggregations weighed significantly more than those on the edges of aggregations (Fig S11).

**Figure 2:**
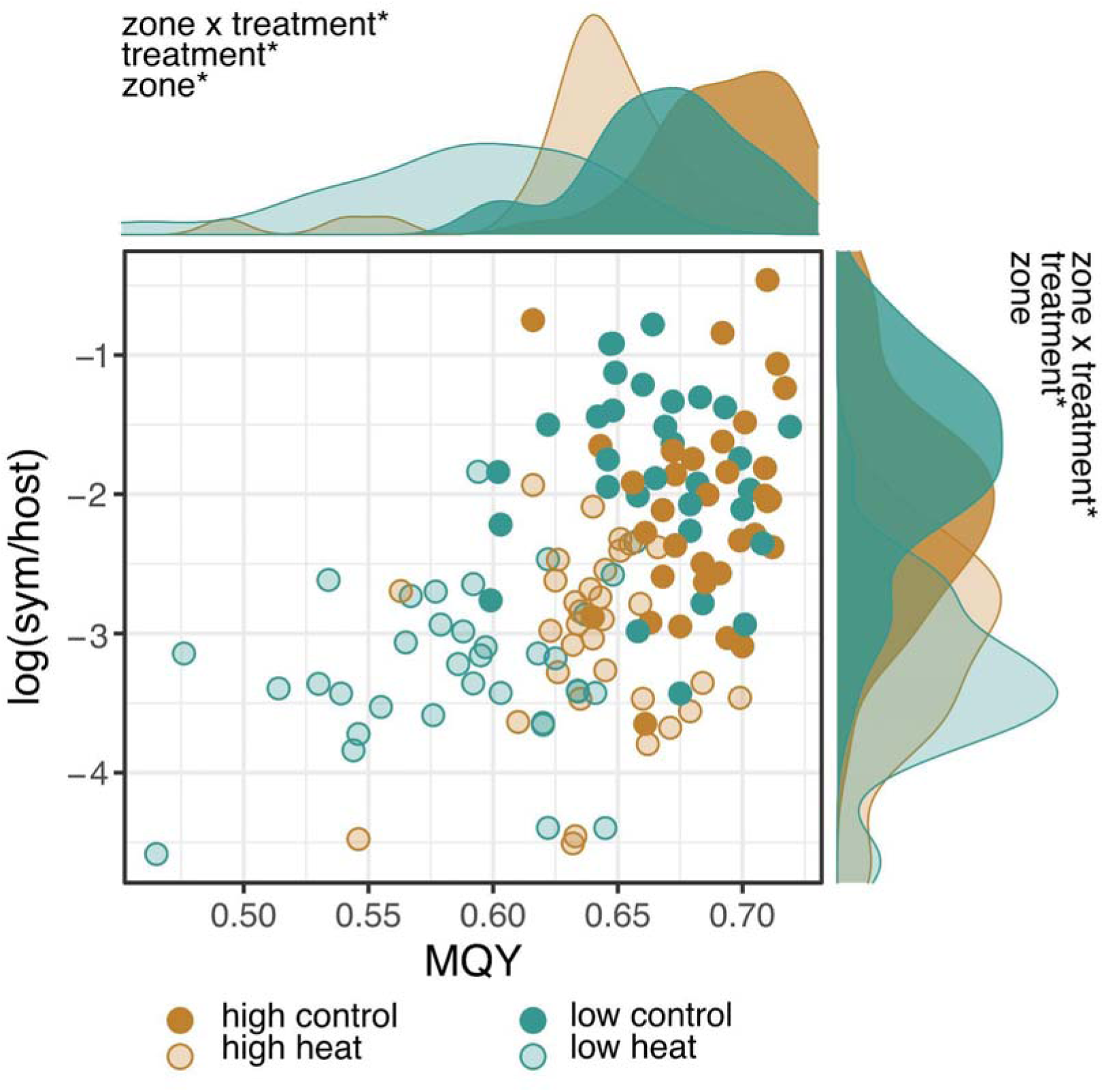
Relationship between Maximum quantum yield (MQY) and symbiont to host cell ratios across tidal zones and treatment. Density plots show the distribution of samples for a given trait (top MQY, right log(sym/host)). Samples from the high intertidal and low intertidal are represented in orange and blue, respectfully. Dark shading represents samples in control conditions whereas light shading represents heat stressed samples. Asterisks (*) denote significance of fixed factors from linear mixed effect models (p < 0.05).

Although independent aggregations were initially assumed to be composed of different host genotypes (McFadden et al., 1997), IBS clustering of 428 SNPs revealed that some aggregations in different zones belonged to the same clonal group (Fig S5). Aggregations 1, 2, 3, 7, and 8 were collapsed into a single genet (Genet 1), as well as aggregations 10 and 11 (Genet 10), resulting in seven, rather than twelve, unique host genotypes. Because clonal aggregations of a single genotype spanned different tidal heights and zones, we were able to explore the effect of microhabitat acclimatization on physiological performance while controlling for host genetic background. Aggregation had a significant effect on MQY (p<0.001) and symbiont to host cell ratios (p<0.001) in Genet 1 (Fig 3, Table S6). However, trait values among aggregations do not perfectly align with tidal height (Fig 3), suggesting additional environmental factors are shifting baselines within clonal groups. Aggregations within Genet 10 reflected population-level trends (Fig 2), where clonal aggregations at elevated tidal heights tended to have higher MQY but lower symbiont to host cell ratios (Fig 3), although these trends were not statistically significant (Table S7). Finally, genotype of the host also affected baseline trait values. Host genet had a significant effect on symbiont to host cell ratios (p<0.01), chlorophyll concentration (p<0.05), and weight (p<0.001), regardless of tidal height or intertidal zone. Genet 1 appears to be driving most of the genotypic effects in symbiont to host cell ratios and chlorophyll, despite significant phenotypic variation among aggregations. Genet 1 had a significantly greater symbiont to host cell ratios than Genet 13 (p<0.01, Fig S12) and marginally higher ratios than Genet 12 (p=0.0536). Genet 1 also had significantly lower chlorophyll concentration compared to Genet 10 (p<0.0203) and marginally less than Genet 4 (p=0.0713, Fig S13). Anemone weight differed significantly across 33% of pairwise comparisons between genets (Fig S14). In contrast, MQY of the symbiont was not affected by host genotype.

**Figure 3:**
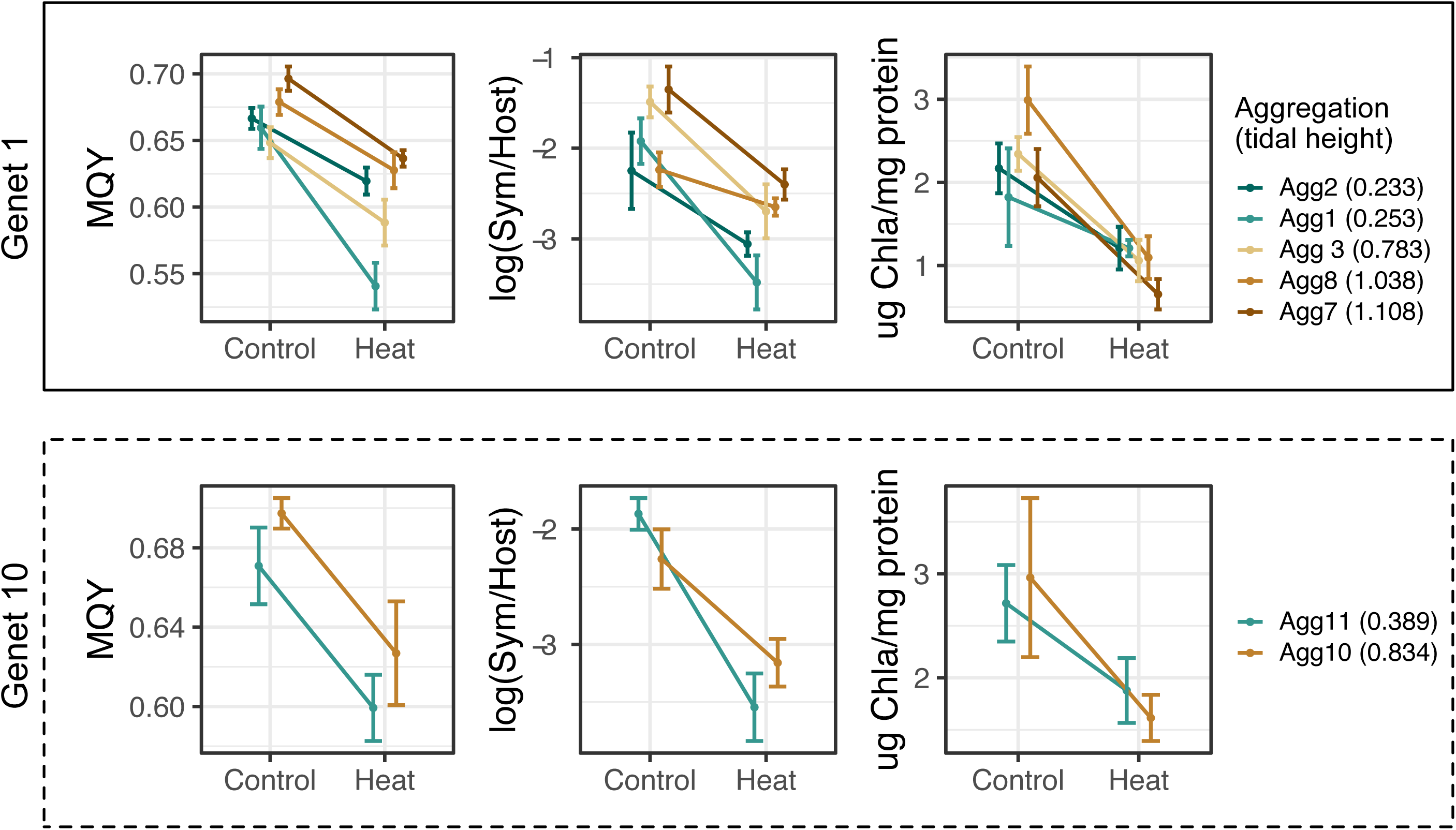
Average trait values +/- SEM (MQY, symbiont to host cell ratios, and chlorophyll a under control and heat stress conditions of clonal aggregations sampled from different tidal heights. Plots are grouped by genotype (top Genet 1, bottom Genet 10). Independent aggregations of the same genet are colored by tidal height. Cooler colors represent aggregations sampled lower in the intertidal and warmer colors represent aggregations higher in the intertidal.

### Response to heat stress

MQY (9%, p<0.001), symbiont to host cell ratios (58%, p<0.001), and chlorophyll a concentration (40%, p<0.001) all declined in anemones exposed to heat stress. There was a significant treatment by intertidal zone interaction for MQY (p<0.05), indicating that symbionts responded differently to heat stress based on intertidal origin. Symbionts from high intertidal anemones had higher baseline MQY which remained elevated under thermal stress compared to symbionts from low intertidal anemones (Fig 2, Fig S15). MQY of high intertidal anemone symbionts declined by 7% on average in heat stress conditions versus a 12% decline in low intertidal anemone symbionts. There was also a significant treatment by intertidal zone interaction on symbiont to host cell ratios (p<0.05). Low intertidal anemones had higher symbiont to host cell ratios under control conditions but ratios declined more under heat stress (71%) compared to high intertidal anemones (46%, Fig 2). Chlorophyll a concentration declined by 40% under heat stress on average, with a reduction of 45% in high intertidal anemones compared to 35% in low intertidal anemones, but this effect was not statistically significant (Fig S15).

No fixed effect of host genotype was observed for the response to heat stress in any physiological trait. However, among aggregations within a genet, microhabitat acclimatization had a variable effect on the response to heat stress. Genet 10’s response reflects the additive zone-level patterns (Fig 2) where aggregations positioned higher in the intertidal (0.834m) maintained about 5% higher MQY values and 11% higher symbiont to host cell ratios on average under heat stress compared to ramets from lower intertidal origin (0.389m), although these differences were not statistically significant (Fig 3). For Genet 1, aggregation and treatment did have a significant interactive effect on MQY (p<0.05) but the response was not driven by tidal height or zone. Under heat stress, MQY was significantly lower for symbionts acclimatized to 0.253m compared to symbionts acclimatized to 1.038m and 1.108m (p<0.001, Fig 3), consistent with results for Genet 10 and the full dataset. However, MQY was also significantly different between clonemates acclimatized to similar tidal heights (0.233m vs 0.253m) under a common heat stress (p<0.001). In this case, symbionts inhabiting clonemates from the lower tidal height maintained higher MQY values under heat stress (Fig 3). Overall symbiont to host cell ratios were not differentially affected by treatment across aggregations of Genet 1. However, anemones from a tidal height of 1.108m had 31% higher symbiont to host cell ratios after heat stress compared to clonemates sampled from 0.253m (p<0.05), demonstrating that microhabitat acclimatization can affect thermal dysbiosis.

## Discussion

Thermal variability is well known to increase thermal tolerance in intertidal organisms across moderate (shorelines) to large spatial scales (latitudinal) (Brahim et al., 2018; Gaitán-Espitia et al., 2014; Gleason et al., 2018; Willett, 2010), but here we show intraspecific divergence in thermal tolerance traits of two cooperative species can occur within meters. Microhabitat acclimatization affected both baseline differences in symbiotic traits and their sensitivity to thermal stress. Higher intertidal anemones had lower symbiont to host cell ratios, but greater photosynthetic efficiency under control conditions, which was better maintained under heat stress, indicating greater thermal tolerance of high intertidal anemones compared to those lower in the intertidal (Fig 2). Symbiont profiles were consistent among all individuals (Fig S8), indicating community-level variation is not driving differences in performance, though population-level variation could not be resolved. Host genotype contributed to physiological variation, but differing baselines persisted for 17 days in common garden conditions, even in genetically identical individuals (Fig 3), suggesting a long-term effect of environmental acclimatization. Further, there was no effect of host genotype on the response to thermal stress in any physiological trait, indicating that environmental history is the primary driver of phenotypic variation in anemones and their symbionts under thermal stress.

### Acclimatization to more extreme environments increases symbiont photosynthetic efficiency but decreases symbiont to host cell ratios

There was a positive correlation between symbiont performance (MQY) and symbiont to host cell ratios, which was modulated by intertidal zone (Fig 2). Lower intertidal anemones had higher symbiont to host cell ratios, but lower MQY in contrast to the lower symbiont to host cell ratios and greater MQY observed in higher intertidal anemones (Fig 2). This suggests that more extreme environments shift symbiotic baselines toward less dense symbiont communities with greater photosynthetic performance. Consistent with population-level results, MQY was also elevated in clonal aggregations from greater tidal heights (Fig 3), despite no change in host genetic background or symbiont community profile, suggesting photosynthetic performance is the result of acclimatization. However, there is some indication that genotypic differences can modulate physiological acclimatization. Genet 10’s symbiont to host cell ratios reflected population-level patterns, but high intertidal aggregations from Genet 1 tended to have greater symbiont to host cell ratios than low intertidal aggregations, opposite of population-level patterns (Fig 3). Host genotype also had a significant effect on symbiont to host cell ratios, but not MQY, suggesting baseline symbiont to host cell ratios may be driven by a combination of genetic and environmental effects.

Symbioses are known to be environmentally responsive with acclimatization reported in several systems, including cnidarian-algal (Putnam, 2021), *Wolbachia*-insect (Mouton et al., 2007), and plant-mycorrhizal symbioses (Ma et al., 2021). In reef-building corals, seasonal acclimatization induces shifts in many physiological traits in both partners, including symbiont pigmentation and photosynthetic performance, host protein and respiration, and holobiont traits, such as symbiont density and photosynthesis/respiration ratios (Scheufen et al., 2017). Seasonal acclimatization to elevated temperature and irradiance during summer months is associated with greater MQY of coral symbionts (Jurriaans & Hoogenboom, 2020; Scheufen et al., 2017) and reduced symbiont densities (Fagoonee et al., 1999; Fitt et al., 2000; Scheufen et al., 2017), which enhances photosynthetic productivity (Scheufen et al., 2017). This pattern is consistent with acclimatization to higher tidal positions observed in the present study, suggesting that light and/or temperature are also the primary environmental drivers. Although we did not observe differences in chlorophyll concentration by tidal zone, chlorophyll was normalized by host protein, which could also be affected by prior environmental history. In tropical corals, both total protein and chlorophyll are lower in summer months compared to winter (Scheufen et al., 2017). Although we could not measure total protein in this study, chlorophyll and protein may co-vary in anemones acclimatized to different thermal conditions resulting in no detectable difference in this trait across tidal zones.

Thermal stress did not affect the relationship between symbiont to host cell ratios and MQY, but both traits exhibited greater proportional declines in low intertidal anemones (Fig 2), indicating a greater thermal tolerance of anemone holobionts acclimatized to more extreme environments. Small but consistent declines in MQY were observed under thermal stress, with low intertidal symbionts experiencing more photodamage than high intertidal symbionts. Reactive oxygen species (ROS) produced by photodamage to symbionts is a key player in the coral bleaching cascade (Lesser, 1996; Weis, 2008) and may have triggered greater bleaching in low intertidal anemones observed here, despite higher baseline densities (Fig 2). Coral symbionts can acclimate to prior thermal exposure by increasing photoprotective mechanisms, which reduces symbiont photodamage and bleaching under subsequent thermal stress (Middlebrook et al., 2008). As symbiont communities are homogenous across the intertidal, greater thermal performance of high intertidal symbionts could be due to acclimatization to more frequent thermal stress. However, higher symbiont densities in coral also increase bleaching susceptibility (Cunning & Baker, 2012), due to greater cumulative oxidative stress produced by larger symbiont populations. Therefore reduced baseline symbiont densities of high intertidal anemones could also be an acclimatory mechanism of the host that increases bleaching resistance.

### Microhabitat acclimatization drives thermal tolerance variation across fine-spatial scales

Microhabitat acclimatization was the primary driver of thermal tolerance variation across the intertidal. MQY and symbiont to host cell ratios were differentially affected by treatment based on intertidal zone (Fig 2), but not by host genotype. Additionally, within genetically identical hosts and symbionts, clones acclimatized to higher intertidal positions performed better under heat stress (Fig 3). In intertidal limpets, one-day exposure to low tide conditions increased thermal tolerance by up to 6°C (Pasparakis et al., 2016), emphasizing the importance of environmental history on tolerance limits. Although tidal height acclimatization has been implicated in physiological variation in several invertebrates (Gleason et al., 2018; Pasparakis et al., 2016), this is the first study to evaluate plasticity in thermal tolerance across the intertidal while controlling for the influence of genetic effects. The present results support the evolutionary theory that variable environments should select for more plastic individuals (Hendry, 2016; Van Tienderen, 1991), especially over small spatial scales when migration is high (Lenormand, 2002; Richardson et al., 2014; Sultan & Spencer, 2002). High intertidal anemones experienced 9.76°C greater daily thermal variation on average and reached maximum temperatures of 9.6°C above low intertidal anemones, which increased holobiont thermal tolerance under a sustained thermal stress (+10°C for 10 days), indicating some cnidarian-algal symbioses do have the capacity to acclimatize to extreme environmental conditions expected under climate change.

Although there is a clear effect of microhabitat acclimatization on physiological baselines and thermal tolerance, there may be other environmental factors driving variation in addition to emersion duration. Within host genotypes, thermally responsive traits were significantly modulated by aggregation (Fig 2, Table S6). However, physiological trait values did not perfectly align with tidal height measurements within genets (Fig 2), suggesting additional environmental factors may differ between aggregations. The intertidal zone has been previously described as an environmental mosaic (Helmuth et al., 2006), where solar and geothermal heating can drive microscale thermal variation within tidal zones (Denny & Harley, 2006; Marshall et al., 2010). It is therefore possible that thermal stress does not increase linearly with tidal height. As other factors such as light stress can also affect thermal tolerance in symbiotic cnidarians independent of solar heating (Brown et al., 2002), additional fine-scale data are needed to pinpoint which environmental factors are affecting physiological variation at an aggregation-level. Nevertheless, an additive effect of intertidal zone on the physiological response to treatment among all genets (Fig 2), and the tendency for high intertidal acclimatization to elevate thermal tolerance within genets (Fig 3), suggests more extreme environments increase thermal tolerance, but experimental manipulations are necessary to test which environmental factors or combinations thereof, modify thermal tolerance in this system.

Additional population-level variation in the symbionts beyond the detection limits of the sequencing method applied here could also play a role in physiological differences. ITS2 amplicon sequencing is the field standard for characterizing symbiont communities, but the multicopy nature of this marker complicates inter-versus intra-genomic variant calling, preventing population-level analyses (Davies et al., 2022). Using 2bRAD sequencing, Cornwell and Hernandez (2021) found *Anthopleura*-associated *B. muscatinei* symbiont genotypes were correlated to benthic community composition within sites, indicating environmental variation may influence symbiont population structure. As strains of *B. minutum* can differ in baseline maximum quantum yield, chlorophyll concentration, growth rate, and the sensitivity of these traits to thermal stress (Bayliss et al., 2019), symbiont population-level variation could also be driving physiological divergence across the intertidal, but this remains to be tested. Additional genetic variation could also be present in the host due to somatic mutations, which cannot be accurately distinguished from technical errors in the absence of high sequencing coverage (Coorens et al., 2021). Because *A. elegantissima* can reproduce asexually, novel variants generated from somatic mutations can be fixed or lost in newly formed clones and drive phenotypic variation (Reusch et al., 2021). Although these additional levels of genetic variation cannot be resolved in the present study, clonal hosts and their symbionts are nonetheless closely related, supporting acclimatization as the prominent driver of physiological variation.

### Limits to microgeographic acclimatization

Position within an aggregation did not affect symbiotic traits, but did affect anemone weight, suggesting anemone size does not tradeoff with thermal tolerance. Anemones from the center of aggregations weighed significantly more than those on the edge of aggregations (Fig S11), which did not affect trait baselines or thermal sensitivity, despite edge anemones experiencing greater daily thermal variation (Fig 1). Size differences within an aggregation is supported by previous research (Francis, 1976) which additionally found anemones in the center of aggregations are more likely to be reproductive than those on the edge. Consistent trait baselines and thermal sensitivity, despite differences in size, suggests there may not be fitness tradeoffs to thermal tolerance maintenance, but rather to greater cumulative stress. This also suggests that differences in daily range of only 0.5-1°C on average are not enough to induce thermal acclimatization, and more extreme environmental divergence such as between aggregations and tidal zones are necessary to increase thermal tolerance. A meta-analysis on coral bleaching reported that a 1°C increase in daily range could reduce bleaching by a factor of 33 (Safaie et al., 2018). However, this was based on thermal characteristics of contemporary reefs, which reflects both adaptation and acclimatization, and may not accurately project within-generational responses to climate change. Thermal priming of coral holobionts to 3-4°C daily variation has had variable success (Bay & Palumbi, 2015; Dilworth et al., 2021), emphasizing a need to understand the relationship between the scale of environmental variability and tolerance limits.

## Acknowledgements

We thank the USC Wrigley Institute for Environmental Studies for funding this work through the Wrigley Bakus Fellowship, Wrigley Summer Fellowship, and Victoria J. Bertics Fellowship. We’d also like to thank the Wrigley staff for providing support and facilities to run this experiment, Bob Carpenter and Nyssa Silbiger for providing necessary equipment for symbiont photophysiology and tidal height measurements, and Samuel Bedgood, Adib Mustofa, and Jenna Dilworth for field assistance.

## Notes

https://github.com/mruggeri55/AeleHeatStress

https://www.ncbi.nlm.nih.gov/bioproject/?term=PRJNA933708

### Competing Interest Statement

The authors have declared no competing interest.

## References

Bates, D., Mächler, M., Bolker, B., & Walker, S. (2015). Fitting Linear Mixed-Effects Models Using lme4. Journal of Statistical Software, 67, 1–48.

Bayliss, S. L. J., Scott, Z. R., Coffroth, M. A., & terHorst, C. P. (2019). Genetic variation in Breviolum antillogorgium, a coral reef symbiont, in response to temperature and nutrients. Ecology and Evolution, 9(5), 2803–2813.

Bay, R. A., & Palumbi, S. R. (2015). Rapid Acclimation Ability Mediated by Transcriptome Changes in Reef-Building Corals. Genome Biology and Evolution, 7(6), 1602–1612.

Bingham, B. L., Freytes, I., Emery, M., Dimond, J., & Muller-Parker, G. (2011). Aerial exposure and body temperature of the intertidal sea anemoneAnthopleura elegantissima. Invertebrate Biology: A Quarterly Journal of the American Microscopical Society and the Division of Invertebrate Zoology/ASZ, 130(4), 291–301.

Brahim, A., Mustapha, N., & Marshall, D. J. (2018). Non-reversible and Reversible Heat Tolerance Plasticity in Tropical Intertidal Animals: Responding to Habitat Temperature Heterogeneity. Frontiers in Physiology, 9, 1909.

Brown, B., Dunne, R., Goodson, M., & Douglas, A. (2002). Experience shapes the susceptibility of a reef coral to bleaching. Coral Reefs, 21(2), 119–126.

Coorens, T. H. H., Oliver, T. R. W., Sanghvi, R., Sovio, U., Cook, E., Vento-Tormo, R., Haniffa, M., Young, M. D., Rahbari, R., Sebire, N., Campbell, P. J., Charnock-Jones, D. S., Smith, G. C. S., & Behjati, S. (2021). Inherent mosaicism and extensive mutation of human placentas. Nature, 592(7852), 80–85.

Cornwell, B. H., & Hernández, L. (2021). Genetic structure in the endosymbiont Breviolum “muscatinei” is correlated with geographical location, environment and host species. Proceedings of the Royal Society B: Biological Sciences, 288(1946), 20202896.

Cunning, R., & Baker, A. C. (2012). Excess algal symbionts increase the susceptibility of reef corals to bleaching. Nature Climate Change, 3(3), 259–262.

Davies, S., Gamache, M. H., Howe-Kerr, L. I., Kriefall, N. G., Baker, A. C., Banaszak, A. T., Bay, L. K., Bellantuono, A. J., Bhattacharya, D., Chan, C. X., & Others. (2022). *Building consensus around the assessment and interpretation of Symbiodiniaceae diversity*. https://repository.kaust.edu.sa/handle/10754/679256

Denny, M. W., & Harley, C. D. G. (2006). Hot limpets: predicting body temperature in a conductance-mediated thermal system. The Journal of Experimental Biology, 209(Pt 13), 2409–2419.

Dilworth, J., Caruso, C., Kahkejian, V. A., Baker, A. C., & Drury, C. (2021). Host genotype and stable differences in algal symbiont communities explain patterns of thermal stress response of Montipora capitata following thermal pre-exposure and across multiple bleaching events. Coral Reefs, 40(1), 151–163.

Dimond, J. L., Bingham, B. L., Muller-Parker, G., Wuesthoff, K., & Francis, L. (2011). Seasonal stability of a flexible algal-cnidarian symbiosis in a highly variable temperate environment. Limnology and Oceanography, 56(6), 2233–2242.

Elder, H. L. (2020). *Genomic Resource Development and Studies of Thermal Tolerance in Symbiotic Cnidarians*. https://ir.library.oregonstate.edu/concern/graduate_thesis_or_dissertations/rr172450q

Fagoonee, I., I., Wilson, H. B., Hassell, M. P., & Turner, J. R. (1999). The dynamics of zooxanthellae populations: A long-term study in the field. Science, 283(5403), 843–845.

Felsenstein, J. (1976). The theoretical population genetics of variable selection and migration. Annual Review of Genetics, 10, 253–280.

Fitt, W. K., McFarland, F. K., Warner, M. E., & Chilcoat, G. C. (2000). Seasonal patterns of tissue biomass and densities of symbiotic dinoflagellates in reef corals and relation to coral bleaching. Limnology and Oceanography, 45(3), 677–685.

Francis, L. (1976). SOCIAL ORGANIZATION WITHIN CLONES OF THE SEA ANEMONE ANTHOPLEURA ELEGANTISSIMA. The Biological Bulletin, 150(3), 361–376.

Gaitán-Espitia, J. D., Bacigalupe, L. D., Opitz, T., Lagos, N. A., Timmermann, T., & Lardies, M. A. (2014). Geographic variation in thermal physiological performance of the intertidal crab Petrolisthes violaceus along a latitudinal gradient. The Journal of Experimental Biology, 217(Pt 24), 4379–4386.

Gleason, L. U., Strand, E. L., Hizon, B. J., & Dowd, W. W. (2018). Plasticity of thermal tolerance and its relationship with growth rate in juvenile mussels (Mytilus californianus). Proceedings. Biological Sciences / The Royal Society, 285(1877). https://doi.org/10.1098/rspb.2017.2617

Helmuth, B., Broitman, B. R., Blanchette, C. A., Gilman, S., Halpin, P., Harley, C. D. G., O’Donnell, M. J., Hofmann, G. E., Menge, B., & Strickland, D. (2006). Mosaic patterns of thermal stress in the rocky intertidal zone: Implications for climate change. Ecological Monographs, 76(4), 461–479.

Hendry, A. P. (2016). Key Questions on the Role of Phenotypic Plasticity in Eco-Evolutionary Dynamics. The Journal of Heredity, 107(1), 25–41.

Hendry, A. P., Day, T., & Taylor, E. B. (2001). Population mixing and the adaptive divergence of quantitative traits in discrete populations: a theoretical framework for empirical tests. Evolution; International Journal of Organic Evolution, 55(3), 459–466.

Hoegh-Guldberg, O. (1999). Climate change, coral bleaching and the future of the world’s coral reefs. Marine and Freshwater Research, 50(8), 839–866.

Hume, B. C. C., Smith, E. G., Ziegler, M., Warrington, H. J. M., Burt, J. A., LaJeunesse, T. C., Wiedenmann, J., & Voolstra, C. R. (2019). SymPortal: A novel analytical framework and platform for coral algal symbiont next-generation sequencing ITS2 profiling. Molecular Ecology Resources, 19(4), 1063–1080.

Jurriaans, S., & Hoogenboom, M. O. (2020). Seasonal acclimation of thermal performance in two species of reef-building corals. Marine Ecology Progress Series, 635, 55–70.

Kelly, M. (2019). Adaptation to climate change through genetic accommodation and assimilation of plastic phenotypes. Philosophical Transactions of the Royal Society of London. Series B, Biological Sciences, 374(1768), 20180176.

Kumarathunge, D. P., Medlyn, B. E., Drake, J. E., Tjoelker, M. G., Aspinwall, M. J., Battaglia, M., Cano, F. J., Carter, K. R., Cavaleri, M. A., Cernusak, L. A., Chambers, J. Q., Crous, K. Y., De Kauwe, M. G., Dillaway, D. N., Dreyer, E., Ellsworth, D. S., Ghannoum, O., Han, Q., Hikosaka, K., … Way, D. A. (2019). Acclimation and adaptation components of the temperature dependence of plant photosynthesis at the global scale. The New Phytologist, 222(2), 768–784.

Lenormand, T. (2002). Gene flow and the limits to natural selection. Trends in Ecology & Evolution, 17(4), 183–189.

Lenth, R. (2022). emmeans: estimated marginal means, aka least-squares means. R package version 1.4. 7. 2020.

Lesser, M. P. (1996). Elevated temperatures and ultraviolet radiation cause oxidative stress and inhibit photosynthesis in ymbiotic dinoflagellates. Limnology and Oceanography, 41(2), 271–283.

Li, A., Li, L., Wang, W., Song, K., & Zhang, G. (2018). Transcriptomics and Fitness Data Reveal Adaptive Plasticity of Thermal Tolerance in Oysters Inhabiting Different Tidal Zones. Frontiers in Physiology, 9, 825.

Manzello, D. P., Matz, M. V., Enochs, I. C., Valentino, L., Carlton, R. D., Kolodziej, G., Serrano, X., Towle, E. K., & Jankulak, M. (2019). Role of host genetics and heat-tolerant algal symbionts in sustaining populations of the endangered coral Orbicella faveolata in the Florida Keys with ocean warming. Global Change Biology, 25(3), 1016–1031.

Marshall, D. J., McQuaid, C. D., & Williams, G. A. (2010). Non-climatic thermal adaptation: implications for species’ responses to climate warming. Biology Letters, 6(5), 669–673.

Ma, X., Geng, Q., Zhang, H., Bian, C., Chen, H. Y. H., Jiang, D., & Xu, X. (2021). Global negative effects of nutrient enrichment on arbuscular mycorrhizal fungi, plant diversity and ecosystem multifunctionality. The New Phytologist, 229(5), 2957–2969.

McFadden, C. S., Grosberg, R. K., Cameron, B. B., Karlton, D. P., & Secord, D. (1997). Genetic relationships within and between clonal and solitary forms of the sea anemone Anthopleura elegantissima revisited: evidence for the existence of two species. Marine Biology, 128(1), 127–139.

Middlebrook, R., Hoegh-Guldberg, O., & Leggat, W. (2008). The effect of thermal history on the susceptibility of reef-building corals to thermal stress. The Journal of Experimental Biology, 211(Pt 7), 1050–1056.

Mouton, L., Henri, H., Charif, D., Boulétreau, M., & Vavre, F. (2007). Interaction between host genotype and environmental conditions affects bacterial density in Wolbachia symbiosis. Biology Letters, 3(2), 210–213.

Palumbi, S. R., Barshis, D. J., Traylor-Knowles, N., & Bay, R. A. (2014). Mechanisms of reef coral resistance to future climate change. Science, 344(6186), 895–898.

Pasparakis, C., Davis, B. E., & Todgham, A. E. (2016). Role of sequential low-tide-period conditions on the thermal physiology of summer and winter laboratory-acclimated fingered limpets, Lottia digitalis. Marine Biology, 163(2), 23.

Putnam, H. M. (2021). Avenues of reef-building coral acclimatization in response to rapid environmental change. The Journal of Experimental Biology, 224(Pt Suppl 1). https://doi.org/10.1242/jeb.239319

Reusch, T. B. H., Baums, I. B., & Werner, B. (2021). Evolution via somatic genetic variation in modular species. Trends in Ecology & Evolution, 36(12), 1083–1092.

Richardson, J. L., Urban, M. C., Bolnick, D. I., & Skelly, D. K. (2014). Microgeographic adaptation and the spatial scale of evolution. Trends in Ecology & Evolution, 29(3), 165–176.

Rosenberg, E., Koren, O., Reshef, L., Efrony, R., & Zilber-Rosenberg, I. (2007). The role of microorganisms in coral health, disease and evolution. Nature Reviews. Microbiology, 5(5), 355–362.

Safaie, A., Silbiger, N. J., McClanahan, T. R., Pawlak, G., Barshis, D. J., Hench, J. L., Rogers, J. S., Williams, G. J., & Davis, K. A. (2018). High frequency temperature variability reduces the risk of coral bleaching. Nature Communications, 9(1), 1671.

Sanders, J. G., & Palumbi, S. R. (2011). Populations of Symbiodinium muscatinei show strong biogeographic structuring in the intertidal anemone Anthopleura elegantissima. The Biological Bulletin, 220(3), 199–208.

Sanford, E., & Kelly, M. W. (2011). Local adaptation in marine invertebrates. Annual Review of Marine Science, 3, 509–535.

Scheufen, T., Krämer, W. E., Iglesias-Prieto, R., & Enríquez, S. (2017). Seasonal variation modulates coral sensibility to heat-stress and explains annual changes in coral productivity. Scientific Reports, 7(1), 4937.

Secord, D., & Augustine, L. (2005). Biogeography and microhabitat variation in temperate algal-invertebrate symbioses: zooxanthellae and zoochlorellae in two Pacific intertidal sea anemones, Anthopleura elegantissima and A. xanthogrammica. Invertebrate Biology: A Quarterly Journal of the American Microscopical Society and the Division of Invertebrate Zoology/ASZ, 119(2), 139–146.

Shoguchi, E., Shinzato, C., Kawashima, T., Gyoja, F., Mungpakdee, S., Koyanagi, R., Takeuchi, T., Hisata, K., Tanaka, M., Fujiwara, M., Hamada, M., Seidi, A., Fujie, M., Usami, T., Goto, H., Yamasaki, S., Arakaki, N., Suzuki, Y., Sugano, S., … Satoh, N. (2013). Draft assembly of the Symbiodinium minutum nuclear genome reveals dinoflagellate gene structure. Current Biology: CB, 23(15), 1399–1408.

Somero, G. N. (2010). The physiology of climate change: how potentials for acclimatization and genetic adaptation will determine “winners” and “losers.” The Journal of Experimental Biology, 213(6), 912–920.

Stillman, J. H., & Somero, G. N. (2000). A comparative analysis of the upper thermal tolerance limits of eastern Pacific porcelain crabs, genus Petrolisthes: influences of latitude, vertical zonation, acclimation, and phylogeny. Physiological and Biochemical Zoology: PBZ, 73(2), 200–208.

Sultan, S. E., & Spencer, H. G. (2002). Metapopulation structure favors plasticity over local adaptation. The American Naturalist, 160(2), 271–283.

Thornhill, D. J., Howells, E. J., Wham, D. C., Steury, T. D., & Santos, S. R. (2017). Population genetics of reef coral endosymbionts (Symbiodinium, Dinophyceae). Molecular Ecology, 26(10), 2640–2659.

Tomanek, L., & Somero, G. N. (1999). Evolutionary and acclimation-induced variation in the heat-shock responses of congeneric marine snails (genus Tegula) from different thermal habitats: implications for limits of thermotolerance and biogeography. The Journal of Experimental Biology, 202(Pt 21), 2925–2936.

Van Tienderen, P. H. (1991). EVOLUTION OF GENERALISTS AND SPECIALISTS IN SPATIALLY HETEROGENEOUS ENVIRONMENTS. Evolution; International Journal of Organic Evolution, 45(6), 1317–1331.

Wang, S., Meyer, E., McKay, J. K., & Matz, M. V. (2012). 2b-RAD: a simple and flexible method for genome-wide genotyping. Nature Methods, 9(8), 808–810.

Weis, V. M. (2008). Cellular mechanisms of Cnidarian bleaching: stress causes the collapse of symbiosis. The Journal of Experimental Biology, 211(Pt 19), 3059–3066.

Willett, C. S. (2010). Potential fitness trade-offs for thermal tolerance in the intertidal copepod Tigriopus californicus. Evolution; International Journal of Organic Evolution, 64(9), 2521–2534.

Wilson, K., Li, Y., Whan, V., Lehnert, S., Byrne, K., Moore, S., Pongsomboon, S., Tassanakajon, A., Rosenberg, G., Ballment, E., Fayazi, Z., Swan, J., Kenway, M., & Benzie, J. (2002). Genetic mapping of the black tiger shrimp Penaeus monodon with amplified fragment length polymorphism. Aquaculture, 204(3), 297–309.

Zilber-Rosenberg, I., & Rosenberg, E. (2008). Role of microorganisms in the evolution of animals and plants: the hologenome theory of evolution. FEMS Microbiology Reviews, 32(5), 723–735.

