## Supplemental figures and tables for "Microhabitat acclimatization shifts physiological baselines and thermal tolerance of the symbiotic anemone, *Anthopleura elegantissima*"

S1. Tidal height of aggregations above or below mean lower low water (MLLW) line. Points represent top, middle, and bottom of aggregations and are colored by intertidal zone designation (low – blue, high – pink).

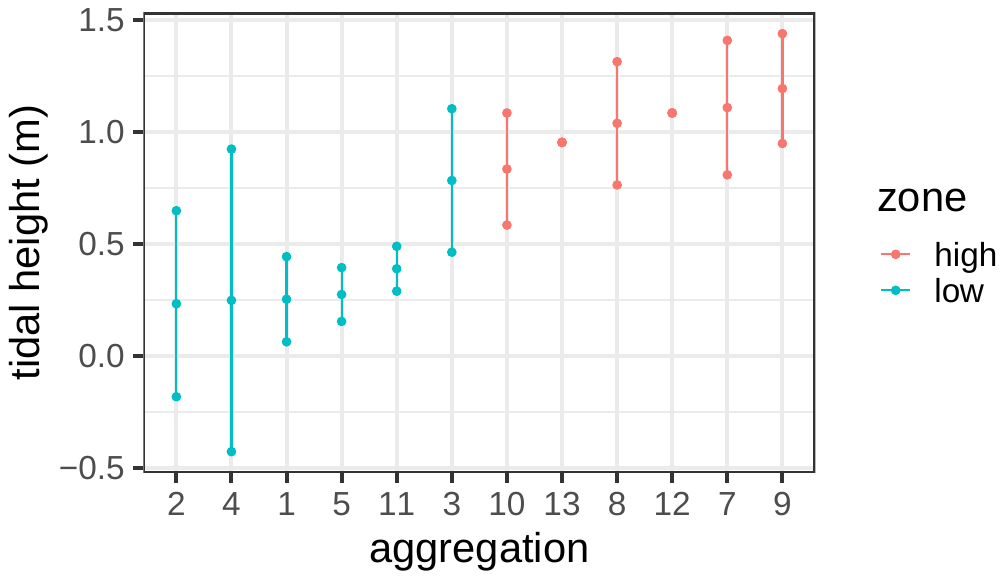

S2. Anemones settled on aragonite plugs in plastic cups to track individuals.

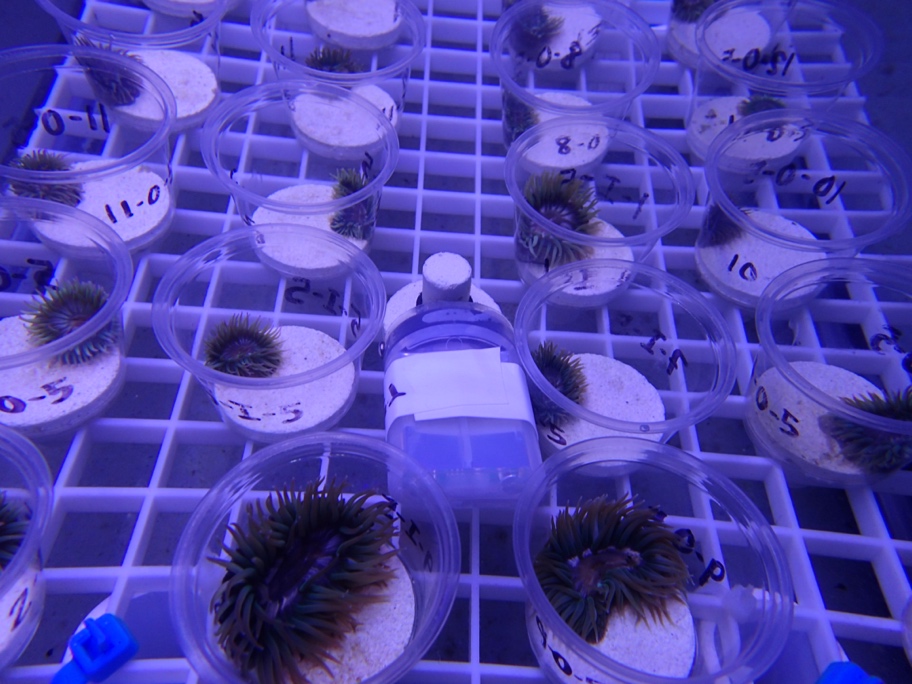

S3. Mean maximum quantum yield (MQY) +/- 1 standard error during the 7-day acclimation period and 10-day heat stress. Dashed line denotes the end of the acclimation period and beginning of experimental heat stress.

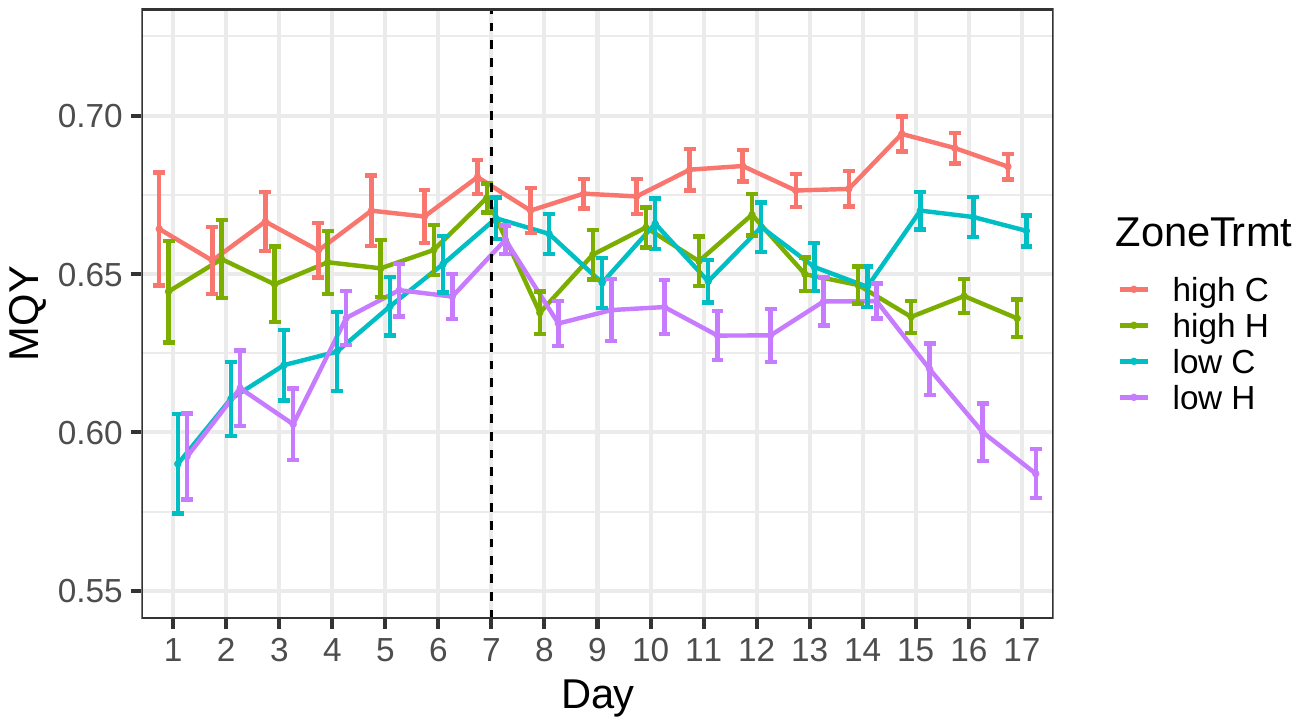

S4. Temperature profiles of three replicate control tanks (pink) and experimental tanks (blue) over the 7-day acclimation period followed by the 10-day experiment.

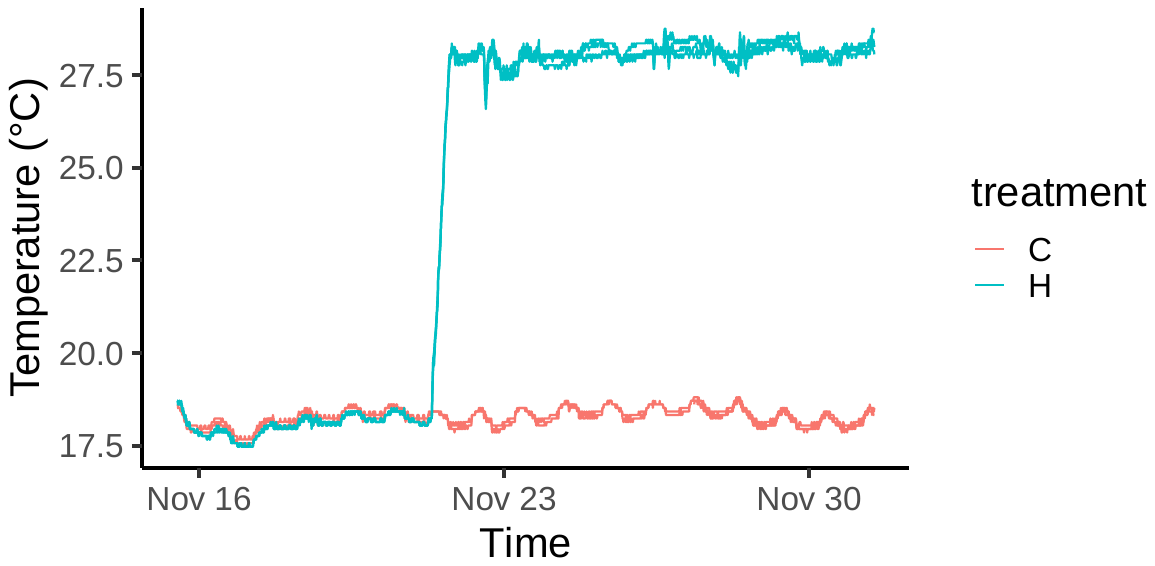

S5. Cluster dendogram of genetic distance (1-IBS) between samples. A subset of four samples per aggregation were sequenced including a representative from each aggregation, position, and treatment combination. Technical replicates colored in red were used to set a distance threshold of 0.22, represented by the dotted line, to distinguish unique genotypes.

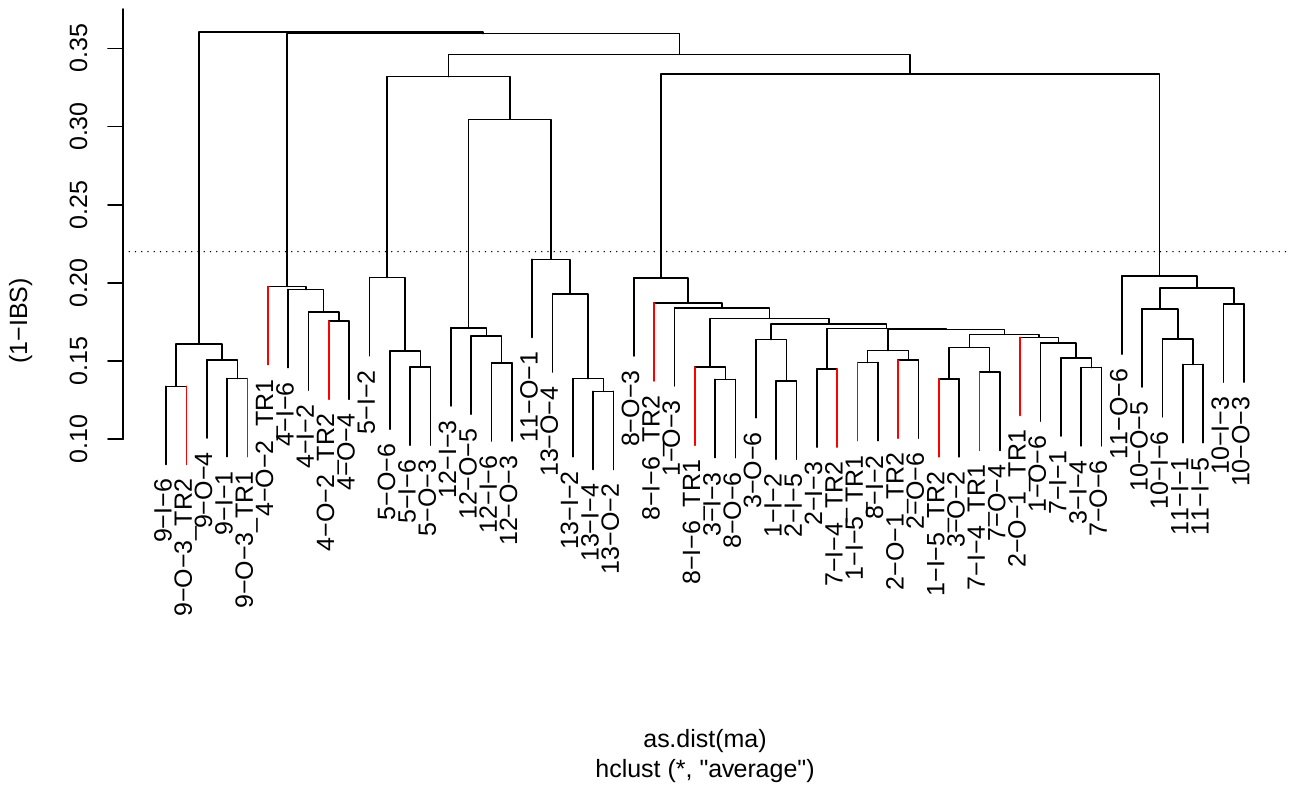

S6. Cluster dendogram of genetic distance (1-IBS) between all replicates of aggregations 11 and 13, confirming that these aggregations are unique genotypes. Technical replicates colored in red were used to set a distance threshold of 0.32, represented by the dotted line.

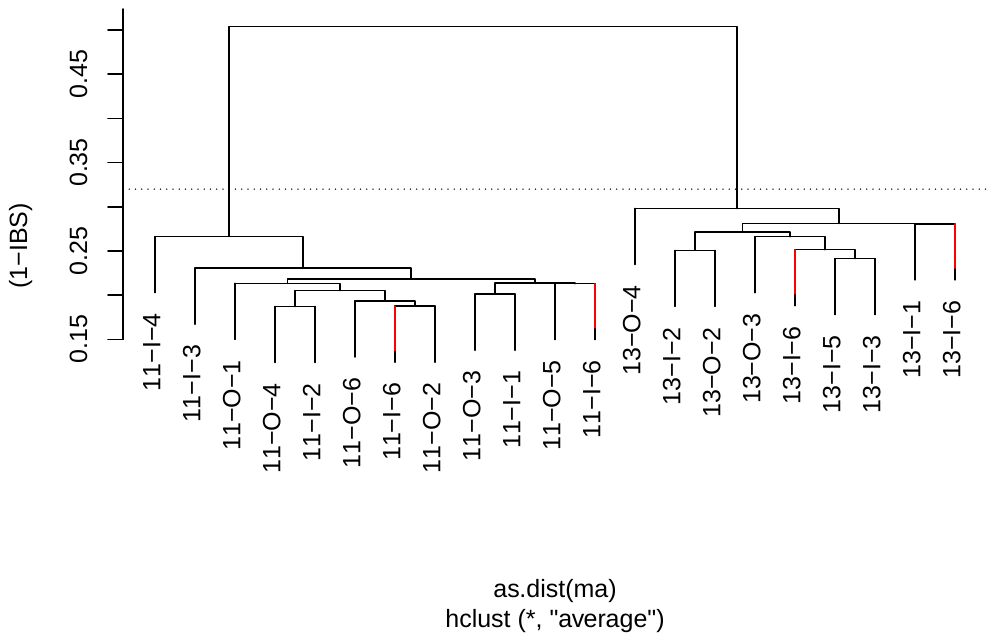

S7. Temperature profile for loggers placed on the center (blue) and edge (pink) of a high intertidal anemone aggregation at an independent site in Los Angeles County, Point Dume.

S8. Relative abundance of ITS2 profiles for each individual. All indivuduals hosted a single profile (1379_B/1393_B-1607_B-B4-1608_B-1418_B-1612_B) corresponding to *Breviolum muscatinei* (blastn 98.51% identity, E=5e^-129^).

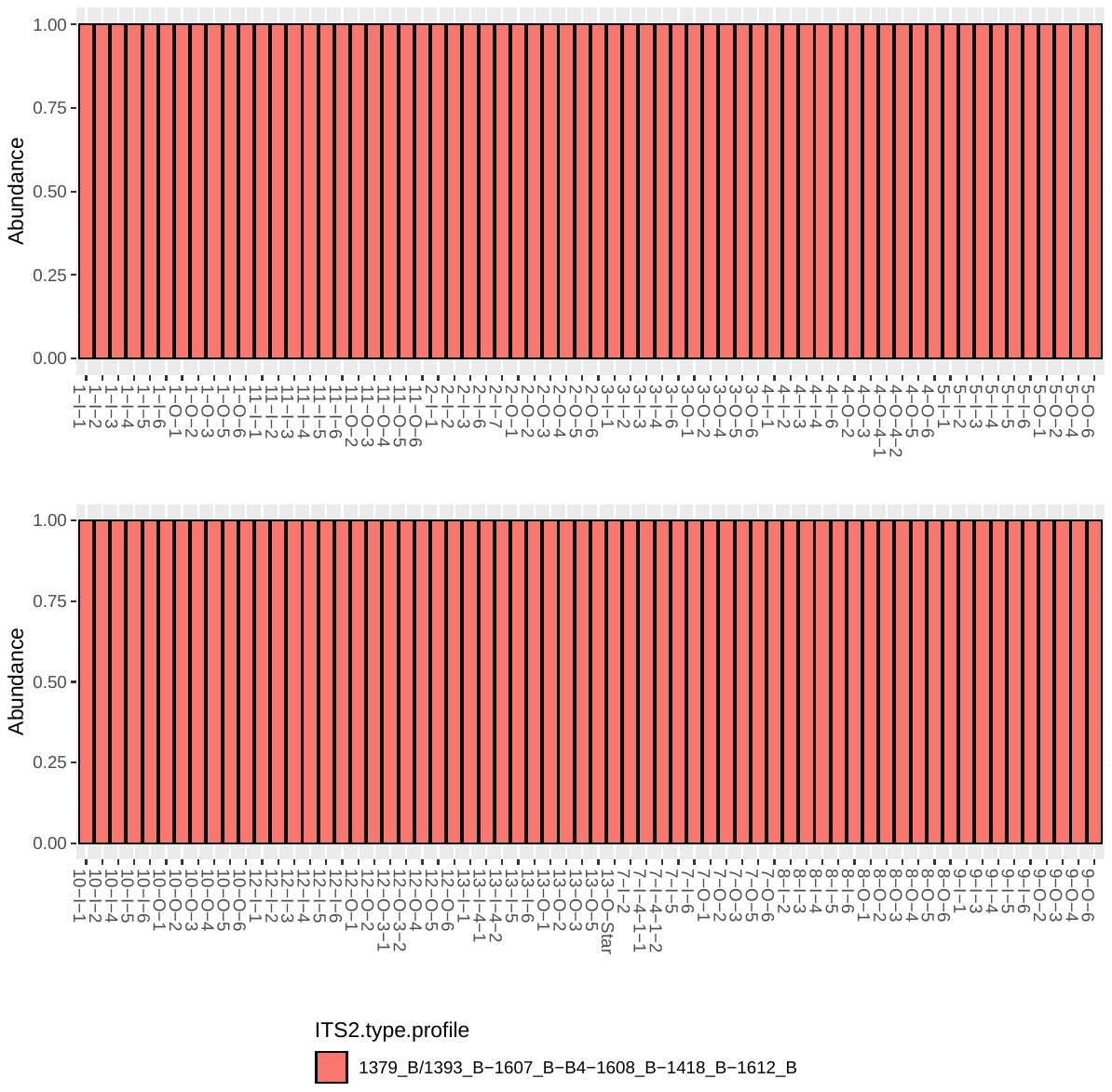

S9. Trait correlations between chlorophyll concentration and MQY (a) and symbiont to host cell ratios (b).

a)	b)

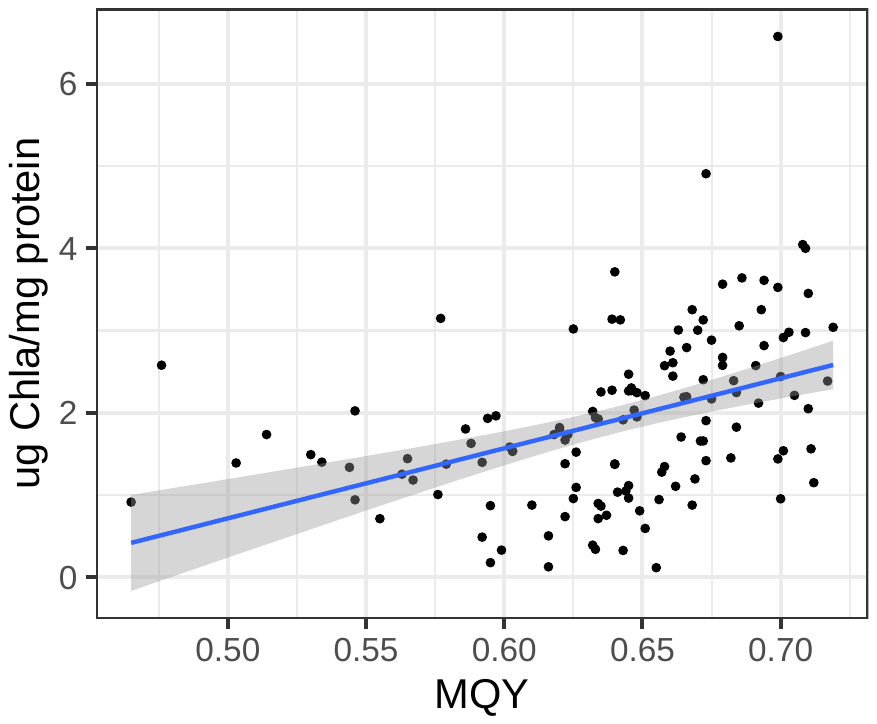

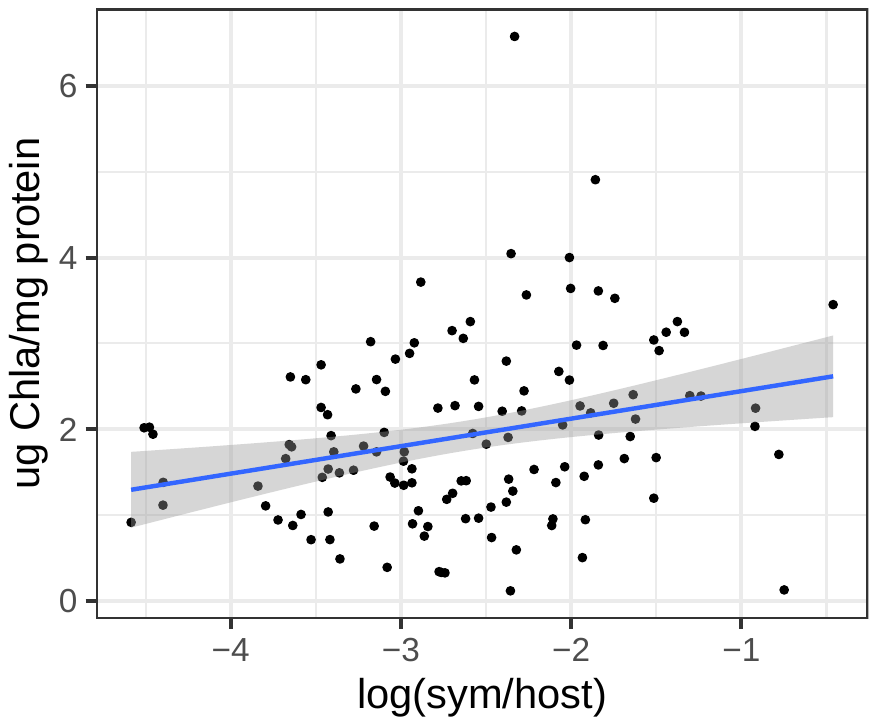

S10. The effect of position within an aggregation (edge – orange, center – blue) on physiological traits.

S11. Initial anemone weight by position within an aggregation (edge vs center).

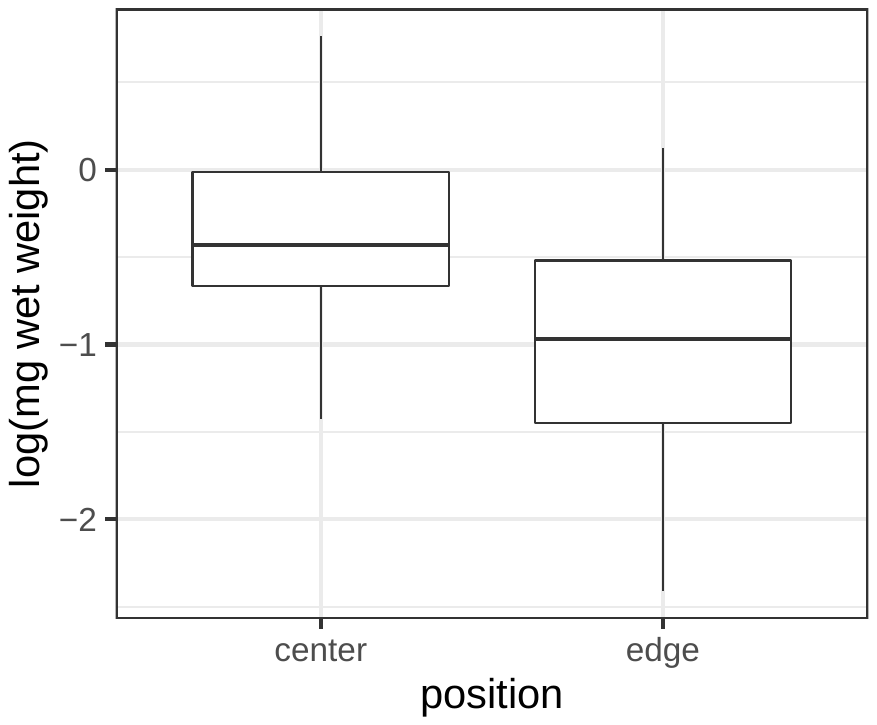

S12. Symbiont to host cell ratios by host genotype.

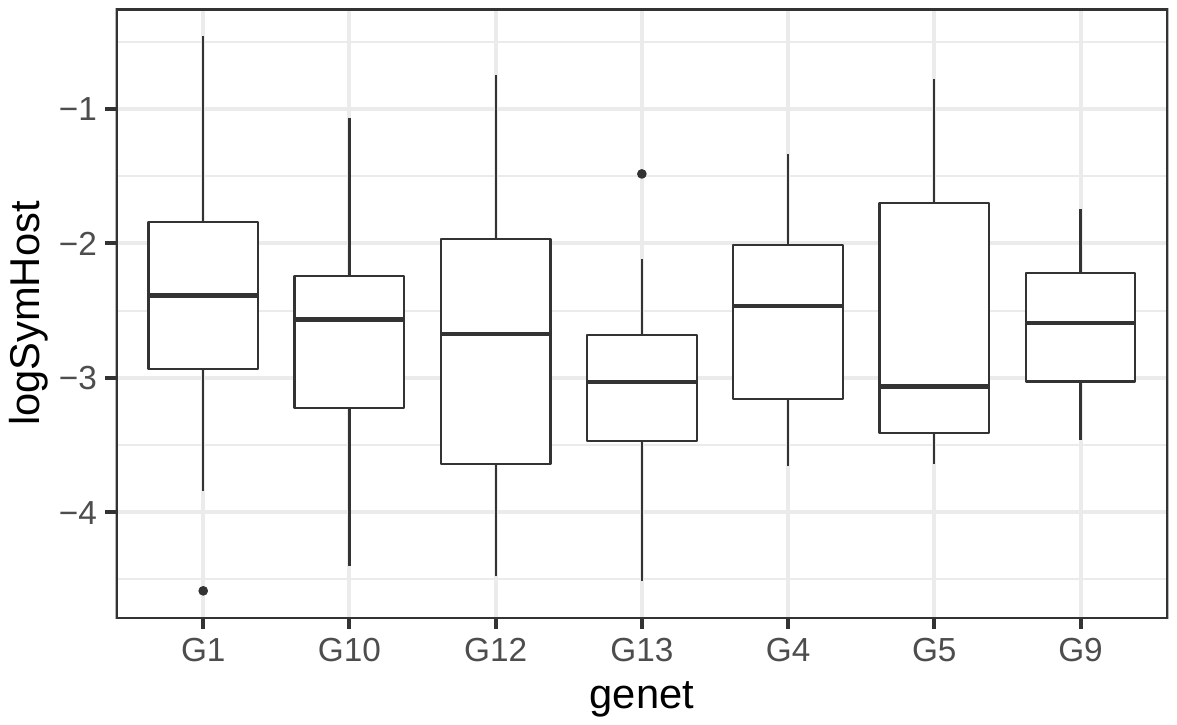

S13. Chlorophyll concentration by host genotype.

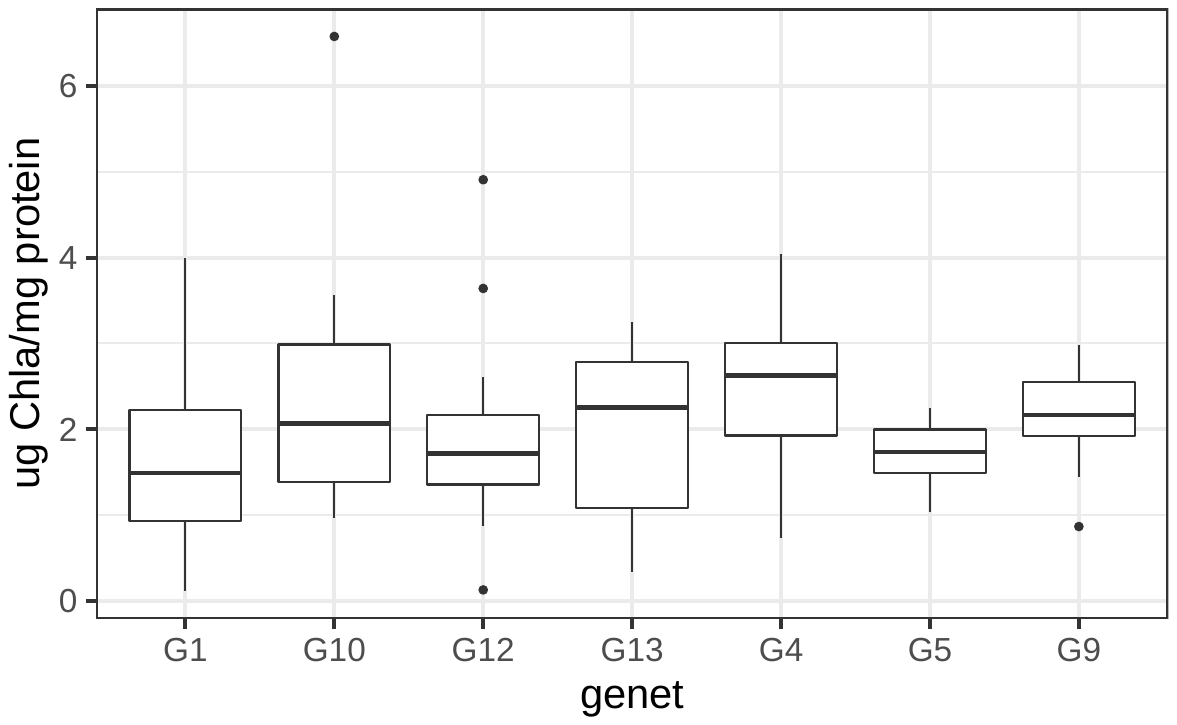

S14. Anemone weight by host genotype.

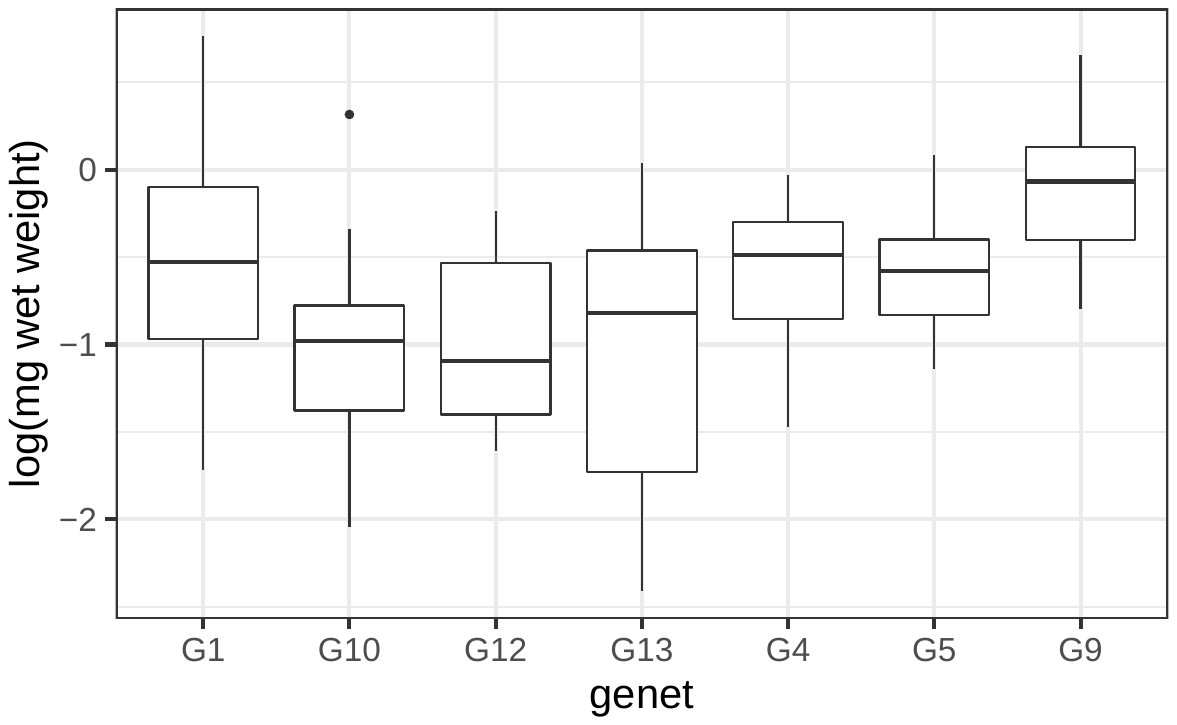

S15. The effect of intertidal zone (high – sand, low – blue) on physiological traits.

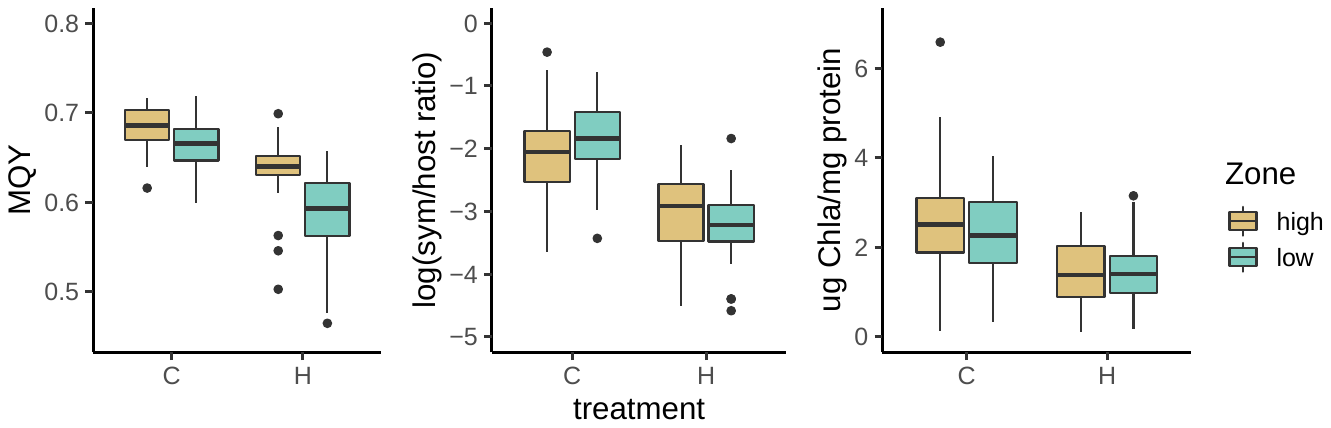

Supplementary tables

S1. Tidal height measurements (m) of bottom, middle, and top of aggregations relative to mean lower low water line.

| **aggregation** | **bottom** | **middle** | **top** |
| --- | --- | --- | --- |
| 1 | 0.631 | 0.821 | 1.011 |
| 2 | 0.386 | 0.801 | 1.216 |
| 3 | 1.031 | 1.351 | 1.671 |
| 4 | 0.141 | 0.816 | 1.491 |
| 7 | 1.376 | 1.676 | 1.976 |
| 8 | 1.331 | 1.606 | 1.881 |
| 9 | 1.516 | 1.761 | 2.006 |
| 13 | 1.521 | 1.521 | 1.521 |
| 5 | 1.19 | 1.31 | 1.43 |
| 10 | 1.62 | 1.87 | 2.12 |
| 11 | 1.325 | 1.425 | 1.525 |
| 12 | 2.12 | 2.12 | 2.12 |

S2. Linear mixed effect model results for all physiological traits (trait ~ treatment*Zone v2 + position + (1|genet) + (1|tank:treatment)) binning aggregation 3 as a high intertidal aggregation. Significant fixed effects below an alpha of 0.05 are bolded.

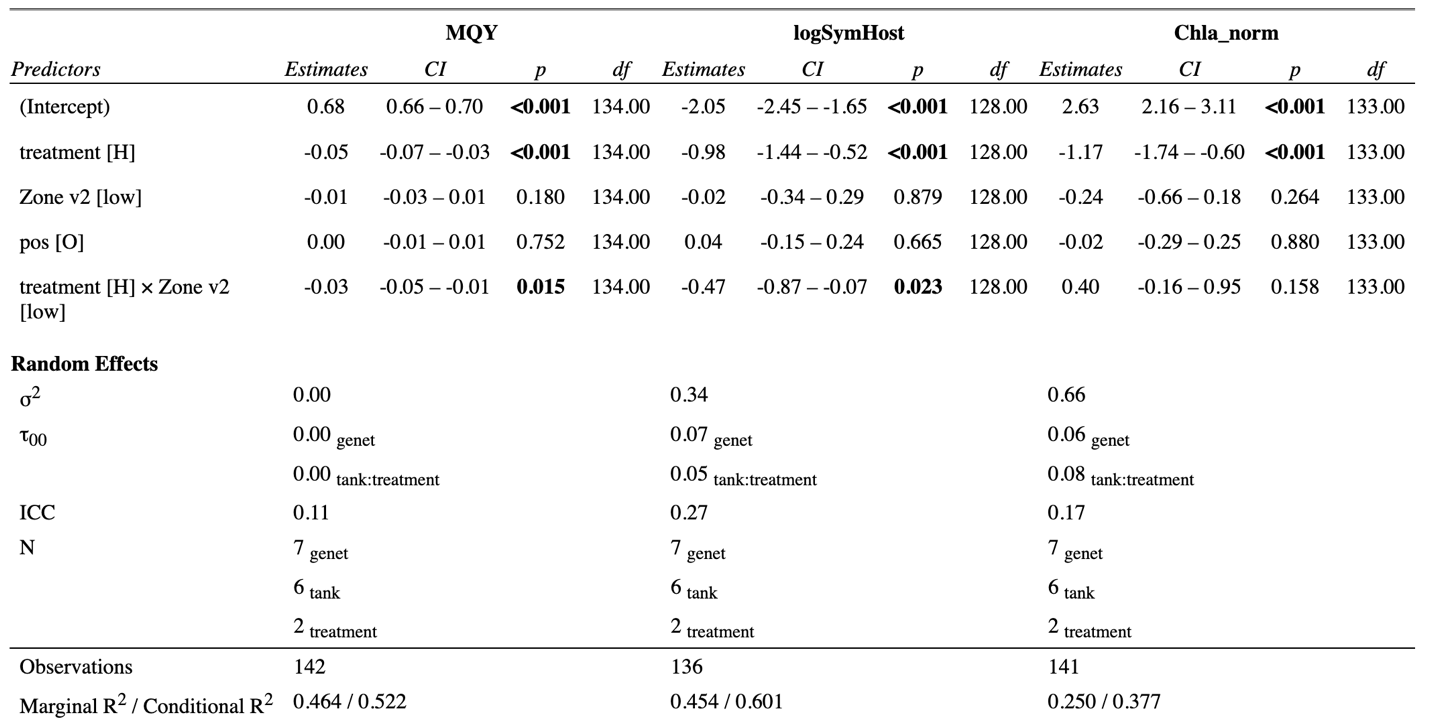

S3. Linear mixed effect model results for all physiological traits (trait ~ treatment*tidal height + position + (1|genet) + (1|tank:treatment)) using tidal height as a continuous variable. Significant fixed effects below an alpha of 0.05 are bolded.

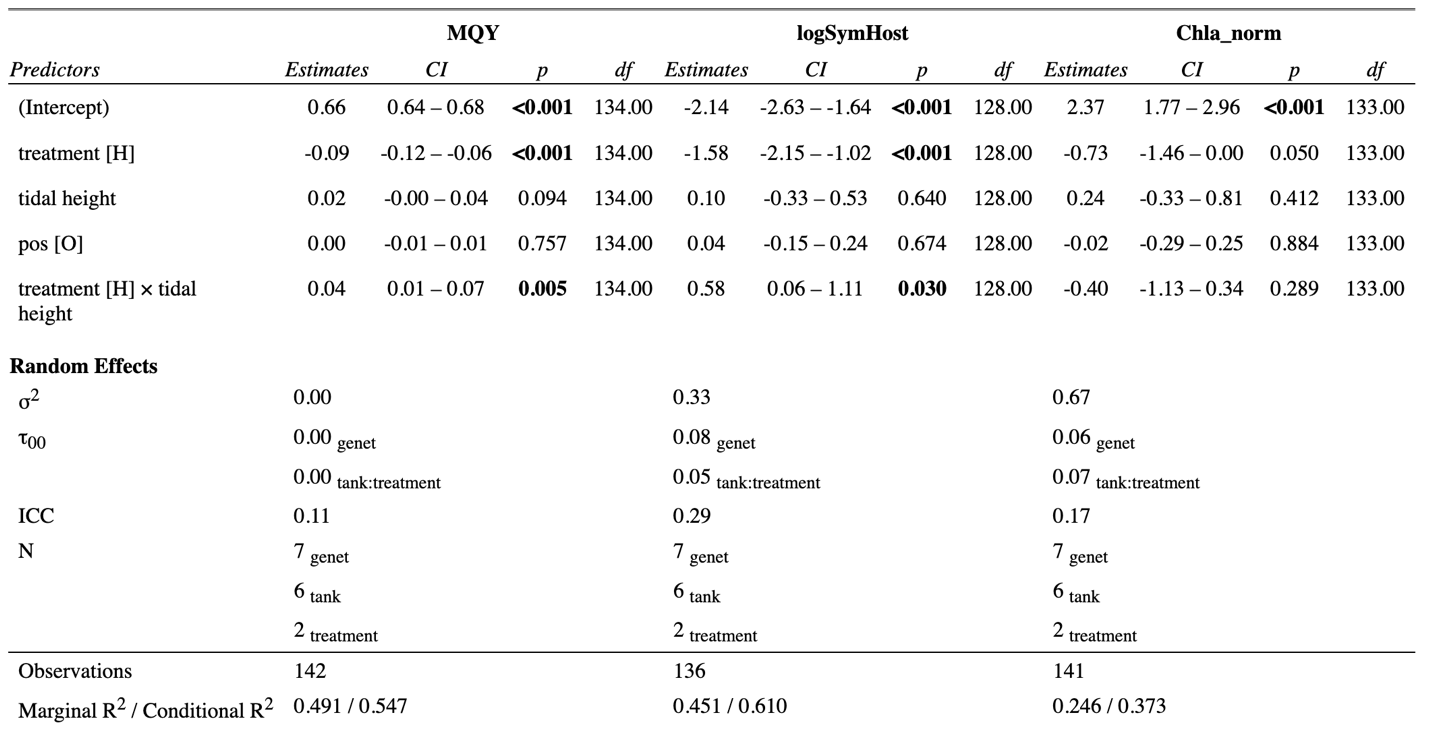

S4. Linear mixed effect model results for all physiological traits (trait ~ treatment*zone + position + (1|genet) + (1|tank:treatment)). Significant fixed effects below an alpha of 0.05 are bolded.

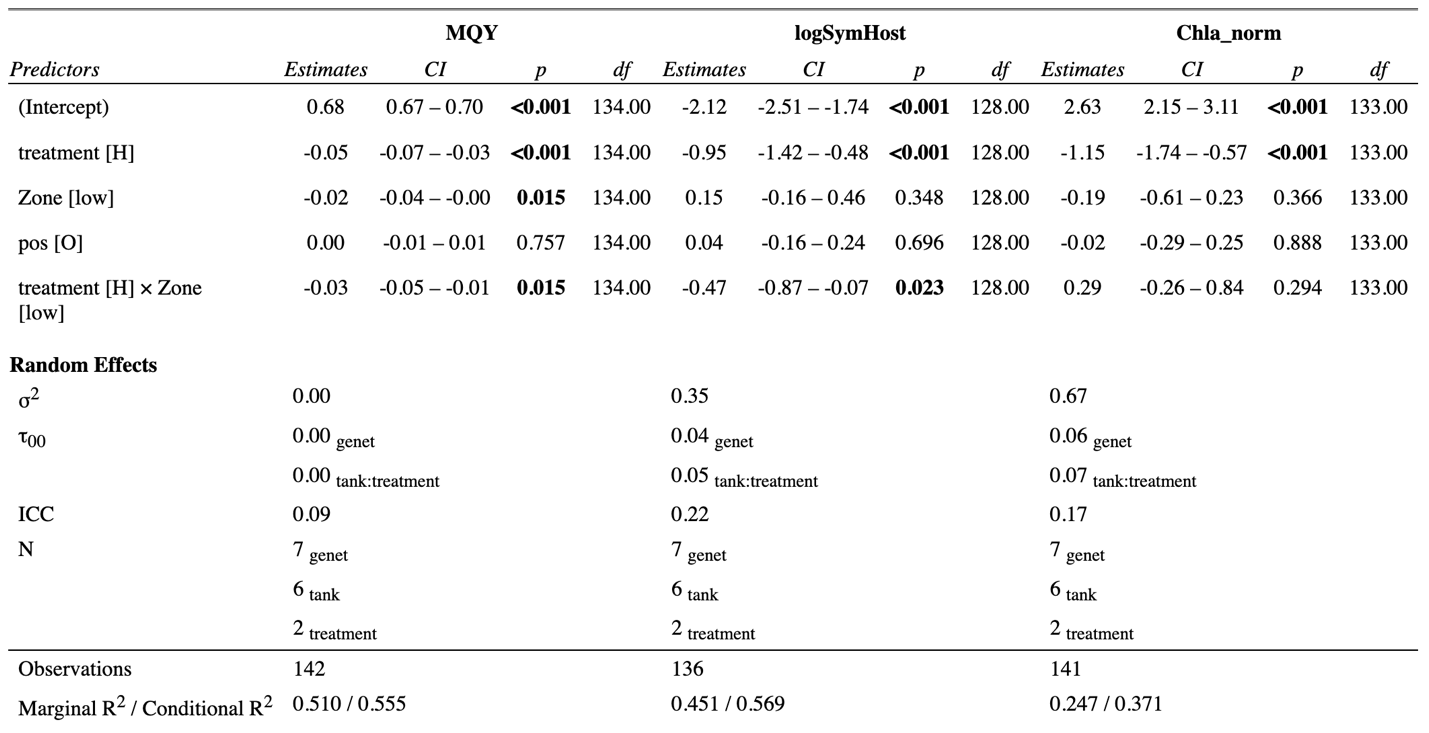

S5. Linear model results for the relationship between symbiont to host cell ratios, maximum quantum yield (MQY), and intertidal zone.

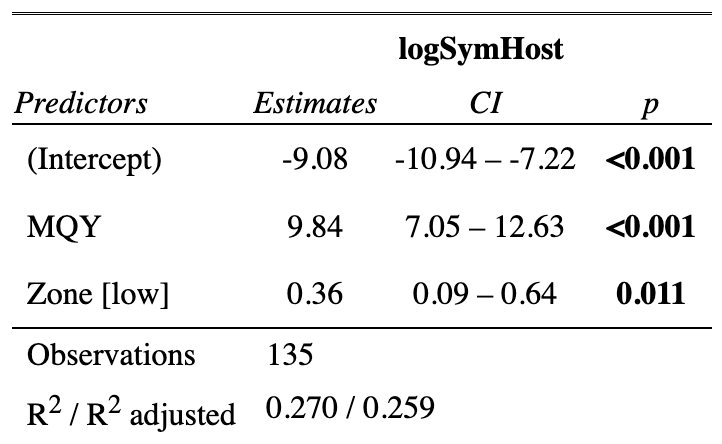

S6. Linear mixed effect model results within Genet 1 for all physiological traits (trait ~ treatment*aggregation + (1|tank:treatment)).

S7. Linear mixed effect model results within Genet 10 for all physiological traits (trait ~ treatment*aggregation + (1|tank:treatment)).
